# Cortical Column and Whole Brain Imaging of Neural Circuits with Molecular Contrast and Nanoscale Resolution

**DOI:** 10.1101/374140

**Authors:** Ruixuan Gao, Shoh M. Asano, Srigokul Upadhyayula, Pisarev Igor, Daniel E. Milkie, Tsung-Li Liu, Singh Ved, Graves Austin, Grace H. Huynh, Yongxin Zhao, John Bogovic, Jennifer Colonell, Carolyn M. Ott, Christopher Zugates, Susan Tappan, Alfredo Rodriguez, Kishore R. Mosaliganti, Sean G. Megason, Jennifer Lippincott-Schwartz, Adam Hantman, Gerald M. Rubin, Tom Kirchhausen, Stephan Saalfeld, Yoshinori Aso, Edward S. Boyden, Eric Betzig

## Abstract

Optical and electron microscopy have made tremendous inroads in understanding the complexity of the brain, but the former offers insufficient resolution to reveal subcellular details and the latter lacks the throughput and molecular contrast to visualize specific molecular constituents over mm-scale or larger dimensions. We combined expansion microscopy and lattice light sheet microscopy to image the nanoscale spatial relationships between proteins across the thickness of the mouse cortex or the entire *Drosophila* brain, including synaptic proteins at dendritic spines, myelination along axons, and presynaptic densities at dopaminergic neurons in every fly neuropil domain. The technology should enable statistically rich, large scale studies of neural development, sexual dimorphism, degree of stereotypy, and structural correlations to behavior or neural activity, all with molecular contrast.

**One Sentence Summary:** Combined expansion and lattice light sheet microscopy enables high speed, nanoscale molecular imaging of neural circuits over large volumes.

## Main Text

Staring deep into the sky from a mountaintop in the American Southwest on a moonless night instills an emotional appreciation of the vastness of space, where the Milky Way galaxy, 60 billion times more massive than the Sun (*1*), represents but one of an estimated two trillion galaxies in the observable universe (*2*). Yet both our emotions and our intellect are made possible by the human brain, a 1.5 kg organ that, despite its small size, is no less complex and remarkable. There, over 80 billion neurons (*3*) connect through ~7,000 synapses each in a network of immense combinatoric complexity. Collectively, humanity creates a truly vast network containing more synapses than there are stars in the observable universe. Understanding the human mind is perhaps the most audacious undertaking in science today.

Further underscoring this challenge is the knowledge that neural structures span a size continuum over seven orders of magnitude in extent, and are comprised of over 10,000 distinct protein types (*4*) collectively essential to build and maintain neural networks. For well over 100 years microscopy has played a central role in revealing this complexity (*5*, *6*). Electron microscopy (EM) has long been able to image down to the level of individual ion channels and synaptic vesicles (*7*), and today can extend this level of detail across the ~0.03 mm^3^ volume of the brain of the fruit fly *Drosophila melanogaster* (*8*, *9*). However, EM creates a grayscale image where the segmentation of specific subcellular components or the tracing of the complete arborization of specific neurons remains challenging, and where specific proteins can rarely be unambiguously identified. Optical microscopy combined with immunofluorescence, fluorescent proteins, or fluorescence *in situ* hybridization (FISH) enables high sensitivity imaging of specific protein expression patterns in brain tissue (*10*, *11*), brain-wide tracing of sparse neural subsets in flies (*12*, *13*) and mice (*14*), and *in situ* identification of specific cell types (*15*, *16*), but has insufficient resolution for dense neural tracing or the precise localization of specific molecular players within critical subcellular structures such as dendritic spines. Super-resolution (SR) fluorescence microscopy (*17*, *18*) combines nanoscale resolution with protein-specific contrast, but bleaches fluorophores too quickly for large volume imaging and, like EM, would require months to years to image even a single *Drosophila melanogaster* brain (table S1).

Given the vast array of molecular species that contribute to neural communication by many mechanisms in addition to the synaptic connections determined by EM connectomics (*19*), and given that the anatomical circuits for specific tasks can vary significantly between individuals of the same species (*20*, *21*), high resolution 3D imaging with molecular specificity of many thousands of brains may be necessary to yield a comprehensive understanding of the genesis of complex behaviors in any organism. Here we describe a combination of expansion microscopy (ExM) (*22*, *23*), lattice light sheet microscopy (LLSM) (*24*), and terabyte-scale image processing and analysis tools (*25*) that achieves single molecule sensitivity and ~60 × 60 × 90 nm^3^ resolution at volumetric acquisition rates ~700x and 1200x faster than existing high speed SR (*26*) and EM (*9*) methods, respectively (table S1). We demonstrate its utility through multicolor imaging of neural subsets and associated proteins across the thickness of the mouse cortex and the entirety of the *Drosophila* brain, while quantifying nanoscale parameters including dendritic spine morphology, myelination patterns, stereotypic variations in boutons of fly projection neurons, and the number of synapses in each fly neuropil region.

### Combining Expansion and Lattice Light Sheet Microscopy (ExLLSM)

In protein retention expansion microscopy (proExM) (*23*), fluorophore-conjugated antibodies (Abs) and/or fluorescent proteins (FPs) that mark the features of interest within a fixed tissue are chemically linked to an infused polyacrylate gel. After protease digestion of the tissue, the gel can be expanded in water isotropically, creating an enlarged phantom of the tissue that faithfully retains the tissue’s original relative distribution of fluorescent tags (fig. S1, supplementary note 1). This yields an effective resolution given by the original resolution of the imaging microscope divided by the expansion factor. Another advantage of digestion is that lipids, protein fragments, and other optically inhomogeneous organic components that are not anchored to the gel are sufficiently removed such that the expanded gel has a refractive index nearly indistinguishable from water and therefore can be imaged aberration-free to a post-expansion depth of at least 500 μm (fig. S2) using conventional water immersion objectives.

Several challenges emerge when attempting to extend ExM to specimens at the mm scale of the fly brain or a mouse cortical column. First, even a modest fourfold expansion, typical of the examples here (table S2), requires effective voxel dimensions of ~30-50 nm on each side to match the full resolution potential of ExM, or ~20 trillion voxels / mm^3^ / color. This in turn necessitates imaging at speeds on the order of 100 million voxels / sec to complete the acquisition in days rather than weeks or more, as well as an image processing and storage pipeline that can handle such high sustained data rates. Second, photobleaching often extinguishes the fluorescence signal from deeper regions of 3D specimens before they can be imaged – a problem that becomes more severe with thicker specimens, longer imaging durations, and/or the higher illumination intensities needed for faster imaging. Finally, since ExM resolution is proportional to imaging resolution, the latter should be as high as possible within these other constraints, while also striving for near-isotropic resolution, so that neural tracing and quantification of nanoscale structures is not limited by the axis of poorest resolution.

To address these challenges, we turned to LLSM (*24*), which sweeps an ultra-thin (~0.4 μm) sheet of laser light through a specimen and collects the resulting fluorescence from above with a high numerical aperture (NA) objective to image it on a high speed camera (supplementary note 2). Confinement and propagation of excitation light within the detection focal plane permits parallel acquisition of data at rates of 10-100 million voxels/sec at low intensities that minimize photobleaching within the plane and eliminates bleaching in the unilluminated regions above and below. Consequently, large volumes of expanded tissue expressing YFP in a subset of mouse cortical neurons can be imaged with uniform signal from top to bottom (Fig. 1A, left). In contrast, the out-of-focus excitation and high peak power at the multiple foci of a spinning disk confocal microscope (SDCM) photobleach the expanded tissue ~10× faster than LLSM (Fig. 1D), rendering deeper regions completely dark (Fig, 1A and B, center), while the sparse illumination of the SDCM focal array slows volumetric acquisition by ~7× (table S1). Another commercial alternative, Airyscan, efficiently images the fluorescence generated at the excitation focus and uses this information to extend the imaging resolution beyond the diffraction limit (*27*, *28*), but images the expanded tissue ~40× slower (table S1) and with ~20× faster bleaching (Fig. 1D) than LLSM.

**Fig. 1.**
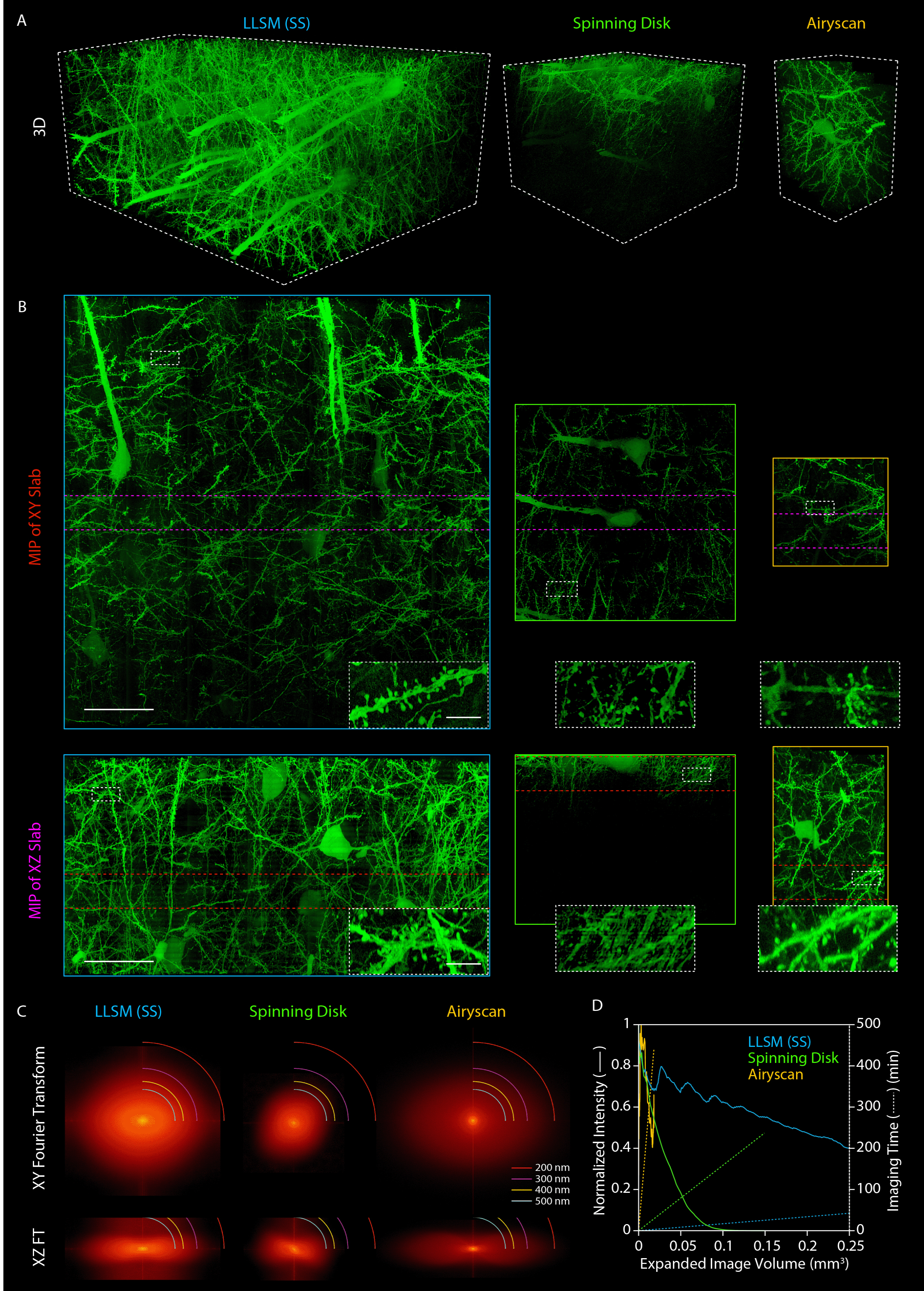
Comparing modalities to image expanded mouse brain tissue. (**A**) 3D rendered volumes at equal magnification of tissue sections from the somatosensory cortex (S1) of a Thy1-YFP transgenic mouse, expanded ~4× using the protein-retention expansion microscopy (proExM) protocol, and imaged by (left to right): lattice-light sheet microscopy in sample scan mode (LLSM (SS), blue); spinning disk confocal microscopy (Spinning Disk, green); and Airyscan in fast mode (Airyscan, orange). (**B**) XY (top) and XZ (bottom) maximum intensity projections (MIPs) of 25 μm thick slabs cut from the image volumes in (A) at the locations denoted by the red and purple lines in the slabs perpendicular to them, respectively. Insets show regions in the white rectangles at higher magnification. Scale bars: 50 μm in full MIPs, 5 μm in inset, here and elsewhere in pre-expanded (i.e., biological) dimensions. (**C**) XY (top) and XZ (bottom) spatial frequency content in the same three image volumes as measured from mitochondria-targeted antibody puncta, with different resolution bands as shown. See also fig. S3. (**D**) Comparative imaging and photobleaching rates for the three modalities. See also table S1.

LLSM can operate in two modes: objective scan (fig. S3), where the sample is stationary while the light sheet and detection objective move in discrete steps across the image volume; and sample scan (Fig. 1), where the sample is swept continuously through the light sheet. Sample scan is faster (tables S1), but yields slightly lower *yz* resolution (fig. S3) than objective scan, since information in the sample scanning direction is slightly blurred by simultaneous image acquisition and sample movement. Of the methods above, Airyscan should in principle achieve the highest lateral (xy) resolution, followed by SDCM (due to pinhole filtering) and finally the two modes of LLSM. In practice, however, dendritic spines and axons appear more clearly and faithfully resolved in lateral views by LLSM than by SDCM or even Airyscan (Fig. 1B, top row), a conclusion corroborated by its higher lateral spatial frequency content (Fig. 1C, fig. S2A, top rows) as measured from mitochondria-targeted Ab puncta. Likewise, the thinness of the lattice light sheet contributes to the axial (z) resolution of LLSM (Fig. 1C, fig. S3A, bottom rows), yielding xz views of spines and axons only slightly poorer than in the lateral plane, and substantially sharper than those obtained by SDCM or Airyscan (Fig. 1B, bottom row).

One additional challenge in mm-scale ExLLSM involves the processing of multi-terabyte (TB) data sets. In LLSM, the lateral extent of the light sheet (table S2) is far smaller than an expanded fly brain or cortical column, so the final image volume must be computationally stitched together from as many as 25,000 (e.g., Fig 6, table S2) tiled subvolumes per color. However, due to systematic sample stage encoder errors and slight swelling or shrinking of expanded samples over many hours, many tiles do not overlap their neighbors on all six sides. To address this, we developed an Apache Spark-based high performance computing pipeline (supplementary note 3, fig. S4-S6) that first performs a flat-field correction for each tile to account for intensity variations across the light sheet and then stitches the intensity-corrected tiles together using an automated and iteratively refined prediction model of tile coordinates. In a separate track, each intensity-corrected tile is deconvolved using a measured point spread function (PSF) so that when the final set of coordinates for all tiles is available, the deconvolved image volume of the entire specimen can be assembled and visualized (supplementary note 4 and 5) with minimal stitching artifacts.

### Quantification of Subcellular Structures in Mouse Cortical Neurons

The protein-specific fluorescence contrast of ExLLSM enables rapid, computationally efficient, and purely automated segmentation and nanoscale quantification of subcellular neural structures over large volumes. For example, dense cytosolic expression of YFP under the *thy1* promotor in mouse pyramidal neurons reveals sharply-delineated voids (Movie 1) representing subcellular compartments (Fig. 2A) of various shapes and sizes whose volumes we could quantify accurately (Fig. 2B, supplementary note 4d). Simultaneous immunofluorescence labeling against Tom20 and LAMP1, although comparatively sparse (movie S1), was sufficient to identify the subset of these that represented mitochondria or lysosomes (Fig. 2C) – in the latter case, the specific subset with LAMP1 that likely represent multivesicular bodies or autolysosomes (supplementary note 6a) (*29*). As expected, we found that mitochondria were generally both longer and larger in volume than lysosomes (Fig. 2D, table S3). Mitochondria ranged in length from 0.2 to 8.0 μm, consistent with EM measurements in the cortex (*30*) or other regions (*31*) of the mouse brain, while the subset of LAMP1 compartments ranged from 0.1 to ~1.0 μm, also consistent with EM (*32*).

**Fig. 2.**
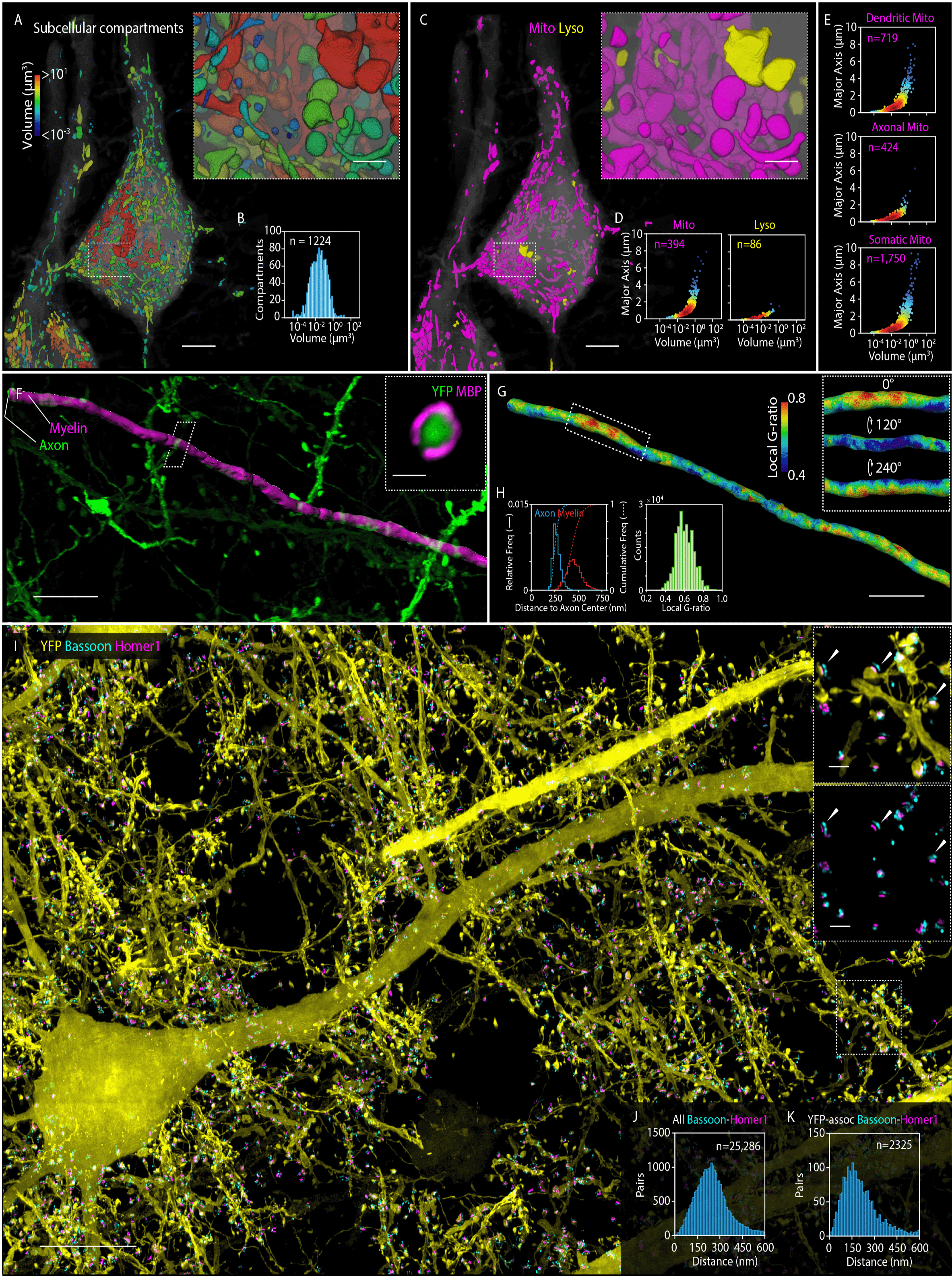
Nanoscale, protein-specific 3D imaging of subcellular neural structures. (**A**) Segmented compartments void of cytosolic YFP (grey), color-coded by volume, in portions of the somata and apical dendrites of two layer V pyramidal neurons from the somatosensory cortex of a Thy1-YFP mouse (Movie 1). Scale bars: 5 μm and 1 μm (inset). (**B**) Distribution of the compartment volumes observed. (**C**) Same region as (A), with voids identified by immunostaining (movie S1) as either mitochondria (magenta) or multivesicular bodies / autolysosomes (yellow). (**D**) Scatter plots of length vs volume for the two organelle types. Point colors in D and E indicate relative data point density (blue: low; red: high). (**E**) Similar scatter plots for mitochondria only, separated by cellular region. See also fig. S7. (**F**) Axon of a layer V pyramidal neuron and its surrounding myelin sheath, from another Thy1-YFP tissue section cut from S1, immunostained against myelin basic protein (MBP, Movie 2), with inset showing a cross-sectional view at a single cut plane (white parallelogram at left). Scale bars: 5 μm and 500 nm (inset). (**G**) Same region as (F), with the myelin sheath color-coded according to the local g-ratio (axon radius / myelin outer radius). Inset shows azimuthal variation in g-ratio in the region within the rectangle at left. Scale bar: 5 μm. See also fig. S9. (**H**) Distribution of axon radius and myelin outer radius (left) and distribution of g-ratio (right) at all points on the axon in (G). **(I**) XY MIP of a 9.3 μm thick slab within a 75 × 100 × 125 μm^3^ volume from layer V of S1 of a Thy1-YFP mouse, immunostained against the pre- and post-synaptic proteins Bassoon and Homer1 (Movie 3). Only Bassoon and Homer1 pairs associated with YFP expressing neurons are shown for clarity. Insets: XY MIPs of a 2.2 μm thick slab from boxed region at right, showing: (top) YFP and YFP-associated Bassoon and Homer1 pairs and; (bottom) all Bassoon and Homer 1 pairs, whether associated with YFP or not. Arrows indicate Bassoon-Homer1 pairs at synapses. See also fig. S10. Scale bars: 10 μm and 1 μm (insets). (**J**) Distribution of distances between paired Bassoon and Homer1 centroids across the entire volume. (**K**) Distribution when restricted to only those pairs associated with YFP expressing neurons.

Given this agreement, and the important roles mitochondria play in dendrite development, synapse formation, calcium regulation, and neurodegenerative disease (*31*, *33*, *34*), we extended our analysis across ~100 × 150 × 150 μm of the mouse somatosensory cortex, classifying length, aspect ratio, and volume (Fig. 2E, fig. S7) of 2893 mitochondria and 222 lysosomes across the somata and initial portions (78 μm mean length) of the apical dendrite of five layer V pyramidal neurons, as well as the initial portions (95 μm mean length) of three descending axon segments. As noted previously in the hippocampus (*33*), we found that long and high aspect ratio mitochondria were far more prevalent in apical dendrites than in axons, with mitochondria longer than 3 μm comprising 6.5% all dendritic mitochondria (~12 per 100 μm of dendrite length) versus 0.7% of all axonal ones. These differences may represent the difficultly in assembling and maintaining large organelles within the narrow confines of the axon, or they may reflect functional differences in the regulation of calcium in axons versus dendrites.

We next turned our attention to the myelination of axons, which is essential for the rapid (*35*, *36*) and energy efficient (*37*) propagation of action potentials (APs) and which, when disrupted, can lead to neurodegenerative diseases such as multiple sclerosis (*38*). The propagation velocity is affected by the g-ratio, the diameter of the axon normalized to the diameter of its surrounding myelin sheath (*39*). Most EM measurements of the g-ratio come from 2D images of single sections cut transversely to axonal tracts (*40*–*42*), and therefore lack information on how the g-ratio might vary along the length of a given axon. To address this, we used ExLLSM to image a 320 × 280 × 60 μm^3^ volume in the primary somatosensory cortex (S1) of a Thy1-YFP transgenic mouse immunostained against myelin basic protein (MBP, Fig. 2F and Movie 2). At every longitudinal position *z* along a given myelinated axon we measured the local g-ratio at every azimuthal position *θ* by dividing the radius *ρ_axon_(θ, z)* of the axon along the radial vector from the axon center by the radius *ρ_myelin_(θ, z)* of the outer edge of the myelin sheath along the same vector (Fig. 2G, fig. S8, supplementary note 4e). Across one 56 μm long segment, the mean g-ratio of 0.57 calculated from mean axon and sheath diameters of 0.52 and 0.90 μm respectively fell at the lower end of a distribution previously reported in the central nervous system, yet was consistent with a theoretical estimate of 0.60 for the ratio that optimizes propagation velocity (*39*). However, these values do not reflect the substantial variability we observed, with the outer axon to outer myelin distance ranging from 0.12-0.35 μm (fig. S9) and the local g-ratio from ~0.4-0.8 (Fig. 2H, Movie 2). Furthermore, the axon and the sheath were rarely concentric, leading to rapid longitudinal changes in capacitance and impedance that may influence the speed and efficiency of signal propagation.

ExLLSM is also well suited to study the nanoscale organization of synaptic proteins over large tissue volumes. Imaging a 75 × 100 × 125 μm tissue section cut from layer IV-V of the primary somatosensory cortex (S1) of a transgenic Thy1-YFP mouse, we identified 25,286 synapses having closely juxtaposed concentrations of immunolabeled pre- and post-synaptic proteins Bassoon and Homer1 (e.g., fig. S10A), 2,325 of which had Homer1 localized at YFP-labeled dendritic spines (Fig. 2I, Movie 3). These tended to form nested caps, with major axis lengths of 856 ± 181 nm and 531 ± 97 nm for Bassoon and Homer1, respectively (median ± median absolute deviation (MAD), fig. S10B,C). The Homer1 distribution is consistent with SR measurements in dissociated hippocampal neurons (DHN) (*43*), but our Bassoon values are slightly larger. The centroid-to-centroid distance we measured between Bassoon/Homer1 pairs was 243 ± 69 nm for all pairs within the volume (Fig. 2J), and 185 ± 70 nm for those associated with YFP-filled spines (Fig. 2K). The difference between these values suggests that mature glutamatergic synapses of layer V pyramidal neurons, which are the ones expressing YFP, are narrower than other types across S1. The difference between these values and previous SR measurements of 150 ± 20 nm in the ventral orbital cortex (n = 252), (*44*), 165 ± 9 in DHN (n = 43) (*43*), and 179 ± 42 nm in the middle of S1 (n = 159) (*45*) may reflect natural variations in different brain regions (*45*) or a systematic bias in these earlier studies arising by measuring the distance between 1D Gaussian fits to the Bassoon/Homer1 distributions in a manually selected slice through the heart of each synapse, versus our approach of calculating the distance between the 3D centroids calculated across the complete distributions.

### Somatosensory Cortex-Spanning Measurement of Dendritic Spines and Excitatory Synapses

The combination of fast imaging (table S1) and targeted sparse labeling enables ExLLSM-based quantification of nanoscale neural structures to be extended to millimeter-scale dimensions over multi-TB data sets. This yields statistically large sample populations that can reveal subtle changes in the distributions of specific morphological parameters across different regions of the brain.

One such application involves the morphology of dendritic spines in different layers of the mouse cerebral cortex. A spine is a small (~0.01-1.0 μm^3^) membranous protrusion from a neuronal dendrite that receives synaptic input from the closely juxtaposed axon of another neuron. Spine morphology has been extensively studied by a variety of imaging methods (*46*), in part because it is related to synaptic strength (*47*), whose time- and activity-dependent change (i.e., plasticity) (*48*) is implicated in learning and memory consolidation (*49*). However, while optical methods such as Golgi impregnations (*50*), array tomography (*10*), confocal (*51*), and two-photon microscopy (*52*, *53*) can image the complete arborization of neurons spanning the cortex, they lack the 3D nanometric resolution needed to measure the detailed morphology of spines. Conversely, EM (*54*, *55*) and SR fluorescence microscopy (*56*, *57*) have the requisite resolution but not the speed to scale readily to cortical dimensions. ExLLSM, however, has both.

To demonstrate, we imaged a 1900 × 280 × 70 μm^3^ tissue slice spanning the pia to the white matter in S1 of a transgenic Thy1-YFP mouse expressing cytosolic fluorescence within a sparse subset of layer V pyramidal neurons, additionally immunostained against Bassoon and Homer1 (Fig. 3A, Movie 4). In each of seven different regions across the cortex (Fig. 3B, fig. S11A), we selected four 27 × 27 × 14 μm^3^ sub-volumes, and used a modified commercial analysis pipeline (supplementary note 4f) (*58*) to segment (fig. S12, movie S2) and measure spine ultrastructure. Across the ~1500 spines so measured, the range of spine head diameters, neck diameters, overall backbone lengths (spine root to tip), and neck backbone lengths (Fig. 3C, fig. S11B, fig. S13) were consistent with those seen in an EM study of layer II/III pyramidal neurons in the mouse visual cortex (*54*), while the absence of spines in the initial segment of the distal apical dendrite, and prevalence of much larger spines on smaller dendritic branches than on the remainder of the distal apical dendrite (Fig. 3D), were in line with an EM study of pyramidal neurons in S1 of the cat (*59*). Indeed, we found that mean spine head diameter and mean neck backbone length each approximately doubled from layer II/III (position 1) to the regions of layers IV and V (positions 3 and 4) nearest the somata, before falling again in layer VI (positions 6 and 7) to levels similar to layer II/III (table S5). This is consistent with a longitudinal *in vivo* study of spine morphology that found that spines closer to the soma, including those on proximal apical dendrites, were more mature and formed stronger synaptic connections than those on basal dendrites or the distal apical dendrite (*60*). We also found that head diameter and backbone length or neck backbone length were correlated across all layers of the cortex (upper columns, Fig. 3C, Fig. S11B, fig. S13, table S4), but neck diameter and neck backbone length were not correlated across all regions (lower columns, Fig. 3C, fig. S11B, table S4).

**Fig. 3.**
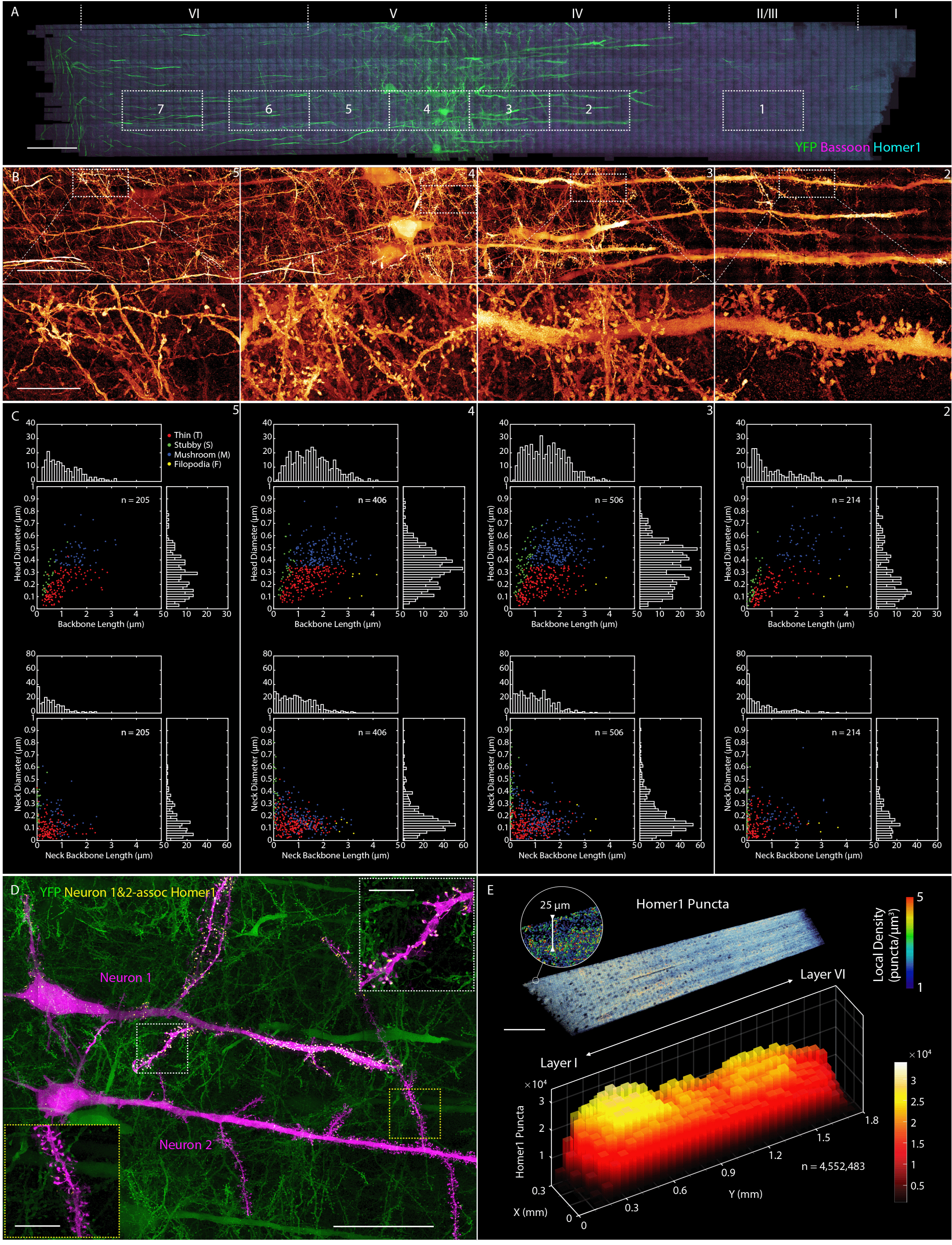
Characterizing dendritic spine morphologies and post-synaptic Homer1 across S1. (**A**) Coronal MIP of a 1900 × 280 × 70 μm^3^ tissue section spanning the pia to the white matter of S1 from a Thy1-YFP mouse (Movie 4), additionally immunostained against Bassoon and Homer1. Boxes denote seven regions for quantitative morphological analysis of dendritic spines. Scale bar: 100 μm. (**B**) Magnified MIPs (top) of YFP expressing neurons in four of the regions from (A), with further magnified sub-regions (bottom) showing differing spine morphologies. Scale bars: 50 μm (top) and 10 μm (bottom). (**C**) Scatter plots and histograms indicating relationships between: spine backbone length and head diameter (top); and spine neck length and neck diameter (bottom) in the four regions from (B). See also movie S2 and figs. S11-S13. (**D**) Two adjacent layer V pyramidal neurons selected within the volume (magenta), one exhibiting strong Homer 1 expression (Neuron 1), and the other exhibiting weak expression (Neuron 2). Insets show Homer1 localization or lack thereof at apical dendritic spines. Scale bars: 50 μm and 10 μm (insets). See also fig. S15. (**E**) MIP of the local density of Homer1 puncta across a ~25 μm thick coronal slab (top), and the cumulative number of puncta in 50 × 50 × 25 μm^3^ subvolumes across the cortex.

Co-labeling with Homer1-specific antibodies allowed us to also map excitatory synapses and their density (Fig. 3E) across S1. In particular, when the 4.5 million Homer1 puncta that we counted were binned in 50 × 50 × 25 μm^3^ sub-volumes to average across local fluctuations, their density was revealed to be ~1.5-2.0× greater in layers II/III and V (~40-50 puncta/μm^3^) than in adjacent layers I, IV and VI, suggesting a greater number of excitatory connections in the former. Similar dual maxima in synaptic density have been observed in sparsely sampled EM images of the rat somatosensory (*61*) and mouse barrel cortex (*62*), although in different cortical layers (II and IV (rat), I and IV (mouse)) than seen here.

Focusing on the subset of Homer1 puncta co-localized with YFP-expressing dendritic spines, we found that thin spines were approximately twice as likely to co-express Homer1 as spines classified as stubby, mushroom, or filopodial (fig. S14). As a synaptic scaffold protein, Homer1 plays an important role in the recruitment and cross-linking of other proteins leading to the maturation and enlargement of spines (*63*–*65*), so its relative abundance at thin spines may presage their transformation to more mature forms. Surprisingly, we also discovered dramatic variations in the expression of Homer1 within neighboring layer V pyramidal neurons, with Homer1 present not only at nearly all spines but also throughout the cytosol of one neuron (Fig. 3D, “Neuron 1”), while a parallel neuron ~57 μm away of similar morphology exhibited very little Homer1, even at its dendritic spines (Fig. 3D, “Neuron 2”). This difference is not the result of differential labeling efficiency, since the density of Homer1 puncta in the immediate surrounds of each neuron is similar (fig. S15). Instead, since Homer1 levels are known to change rapidly under different neuronal states (e.g., asleep vs. awake (*66*)), it may reflect the different excitatory states of these two neurons at the time the animal was sacrificed.

### Visual Cortex-Spanning Neuronal Tracing and Myelination Patterns

While the radial anisotropy of axonal myelination observed in Fig. 2E can affect the speed and efficiency of AP propagation, so too can its longitudinal variation. In fact, the repeated gaps in myelination at the nodes of Ranvier house ion channels that are essential to regenerate the AP during saltatory conduction (*67*), which is the hallmark of high speed signal propagation in vertebrates. Recently, however, high throughput EM imaging and axonal tracing at 30 × 30 × 240 nm^3^ / voxel (*68*) has additionally revealed gaps in the axonal myelination of layer II/III neurons in the mouse visual cortex (V1) much larger (e.g., 55 μm) than either the ~2 μm typical of the nodes of Ranvier or the shorter and rarer gaps observed in layers III to VI of S1 in the same study.

To determine if these differences are more reflective of the layer of origination of the axon or the functional role of the cortical region studied (S1 vs V1), we imaged at 27 × 27 × 50 nm^3^ / voxel a ~280 × 1100 × 83 μm^3^ tissue section from V1 extending from the pia to the white matter of a Thy1-YFP mouse, additionally immunostained against MBP and contactin associated protein (Caspr) (*69*) to visualize myelin sheaths and their terminations, respectively (Fig. 4A, Movie 5). While the dense global staining of EM makes long range 3D tracing of small neurites challenging, expression of YFP in a sparse subset of layer V and layer VI pyramidal neurons (*70*) enabled rapid semi-automatic tracing (supplementary note 4h) of not only axons and their myelination, but also the entire arborization of selected neurons across the entire tissue section (Fig. 4B, Movie 6), including the distal apical dendrite and its branches (Fig. 4C, part i), basal dendrites and their spines (Fig. 4C, part ii), the premyelin axonal segment (PMAS, Fig. 4D), the nodes of Ranvier (Fig. 4E), and collateral branches of the main axon originating at the nodes (Figs. 4F). All these features matched the known morphologies of layer V pyramidal neurons (*71*), and were recapitulated in a second neuron traced throughout the volume (Fig. 4G, Movie 6).

**Fig. 4.**
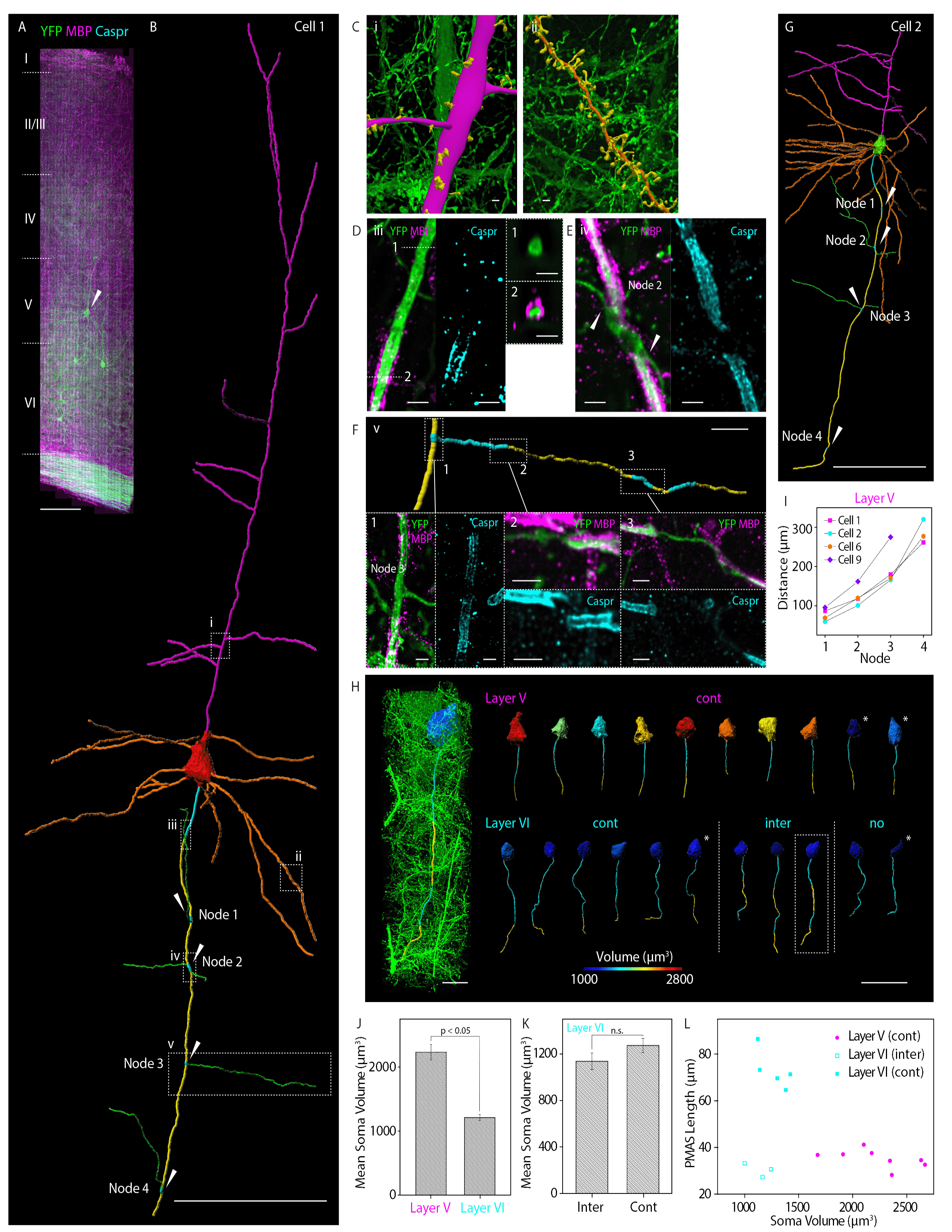
Neural tracing and longitudinal myelination analysis across the primary mouse visual cortex (V1). (**A**) Coronal MIP of a 25 μm thick slab within a 1100 × 280 × 83 μm^3^ tissue section spanning the pia to the white matter of V1 from a Thy1-YFP mouse (Movie 5), additionally immunostained against MBP and Caspr to highlight myelin sheaths and nodes of Ranvier, respectively. Scale bar: 100 μm. (**B**) Traced arborization (Movie 6) of a specific layer V pyramidal neuron denoted by the arrow in (A), showing the soma (red), apical (magenta) and basal (orange) dendrites, myelinated (yellow) and unmyelinated (cyan) axon segments, and collateral axon branches (green). Arrows indicates nodes of Ranvier. Scale bar: 100 μm. (**C**) Magnified segmented views of the distal apical dendrite and two of its branches (left), and a basal dendrite and its spines (right), from boxed regions i and ii in (B), respectively. Scale bars: 1 μm. (**D**) MIP view of boxed region iii in (B), showing: the distal end of the premyelin axonal segment (PMAS) (left); Caspr at the start of myelination (middle); and cross-sectional views of the axon before and after the start of myelination (right). Scale bars: 1 μm. (**E**) MIP view of boxed region iv in (B), showing break in myelination and two branching collateral axons at a node of Ranvier (left), and Caspr highlighting the two ends of the node (right). Scale bars: 1 μm. (**F**) Segmented view (top) of a collateral axon with myelinated and unmyelinated sections from boxed region v in (B), with three MIP views (bottom) of breaks in myelination with flanking Caspr. Scale bars: 10 μm (top) and 1 μm (bottom). (**G**) Traced arborization of a second layer V pyramidal neuron within the volume. Scale bar: 100 μm. (**H**) (Left) Segmented soma and axon of a pyramidal neuron shown in the context of its surroundings in layer VI. (Right) Segmented somata (color-coded by volume) and axons, showing myelinated (yellow) and unmyelinated (cyan) segments, for ten pyramidal neurons from layer V (top row) and eleven more from layer VI (bottom row). Boxed neuron was shown at left. Scale bars: 10 μm (left) and 50 μm (right). (**I**) Node spacing for four layer V neurons from (H). See also fig. S17. (**J**) Volumes of eight layer V and nine layer VI somata fully within the image volume (i.e., no asterisks in (H)) (mean ± SEM). (**K**) Volumes of the three somata with intermittently myelinated axons and five somata with continuously myelinated axons in layer VI (mean ± SEM). The *p* values are calculated from a permutation test for medians (n.s. = not significant). (**L**) Scatter plot of soma volume versus PMAS length for the neurons in (H). See also fig. S16.

Given this assurance, we traced the axons and their longitudinal myelination patterns for 10 neurons in layer V and 11 more in layer VI (Fig. 4H). Within the imaged volume, all of the layer V axons in V1 exhibited continuous myelination beyond the end of the PMAS, except for the expected small gaps at the nodes of Ranvier. This is consistent with the myelination pattern seen previously for layer III to VI axons in S1 (*68*). The range of PMAS lengths we measured for these neurons (28-41 μm, mean = 34.9 ± 1.1 μm) was also consistent with the range found in layers V and VI of S1 (25-40 μm, mean = 33.7 ± 2.4 μm). Notably, the internodal spacing for four of these neurons increased with increasing distance from the soma (Fig. 4I). In contrast, in layer VI only six axons were continuously myelinated, while two were completely unmyelinated, and three exhibited intermittent myelination with long unmyelinated segments more reminiscent of the layer II/III axons in S1 than the layer VI axons there (*68*). Thus, myelination patterns of axons in V1 and S1 can differ, even for neurons in the same cortical layer.

Although the volumes of the somata and the diameters of the PMAS in layer V of V1 were twice as large as those in layer VI (Fig. 4J and fig. S16, respectively), there was not a strong relationship between soma volume and myelination pattern (e.g., intermittent or continuous) within layer VI (Fig. 4K). Notably, however, the PMAS lengths of the six continuously myelinated and the three intermittently myelinated axons in layer VI of V1 split into distinct populations (Fig. 4L), with the intermittent ones of mean length (30.3 ± 1.7 μm) similar to the axons of layer V, and the continuous ones more than twice as long (70.6 ± 3.6 μm). Thus, continuously myelinated axons in different layers of V1 need not have similar PMAS lengths. Given that the distal end of the PMAS is the site of AP initiation (*72*), perhaps PMAS length might be one mechanism by which neurons control the AP to account for differences in myelination or overall axon length in different layers and cortical regions.

### Long-Range Tracing of Clustered Neurons in *Drosophila* and Their Stereotypy

While mm-scale tissue sections present no problem for LLSM, the entire mouse brain is far too large, given the short working distances of commercially available high resolution objectives. The brain of the fruit fly *Drosophila melanogaster*, on the other hand, fits comfortably within the microscope, even in its 4× expanded form. Furthermore, a vast array of genetic tools have been developed for *Drosophila*, such as split-GAL4 drivers and MultiColor FlipOut (MCFO) (*21*), which enable precise labeling of user-selected subsets of its ~100,000 neurons. Fluorescence imaging of thousands of such subsets across thousands of transgenic flies and collation of the results then yields brain-wide 3D reconstructions of complete neural networks at single cell resolution (*12*, *13*). However, to trace fine neuronal processes and identify synaptic connections, nanoscale resolution is needed. For all these reasons, the *Drosophila* brain is well matched to the capabilities of ExLLSM.

To demonstrate, we chose to start with a relatively simple case: three olfactory projection neurons (PNs) originating at the DC3 glomerulus of the antennal lobes that feed most prominent sensory inputs to the calyx (CA) of the mushroom body and lateral horn (LH) (*73*, *74*). Imaging a ~250 × 175 × 125 μm^3^ volume, we were able to trace the axonal branches of all three DC3 PNs across one hemisphere (Fig. 5A, Movie 7), although tracing of fine dendritic processes was still difficult at 4.0× expansion. We were also able to precisely assign boutons to each cell within the CA (Cell 1: 3 boutons, Cell 2: 3 boutons, Cell 3: 4 boutons) and the LH (Cell 1: 19 boutons, Cell 2: 32 boutons, Cell 3: 23 boutons), and determine the shapes and sizes of the boutons in these regions (Fig. 5B).

**Fig. 5.**
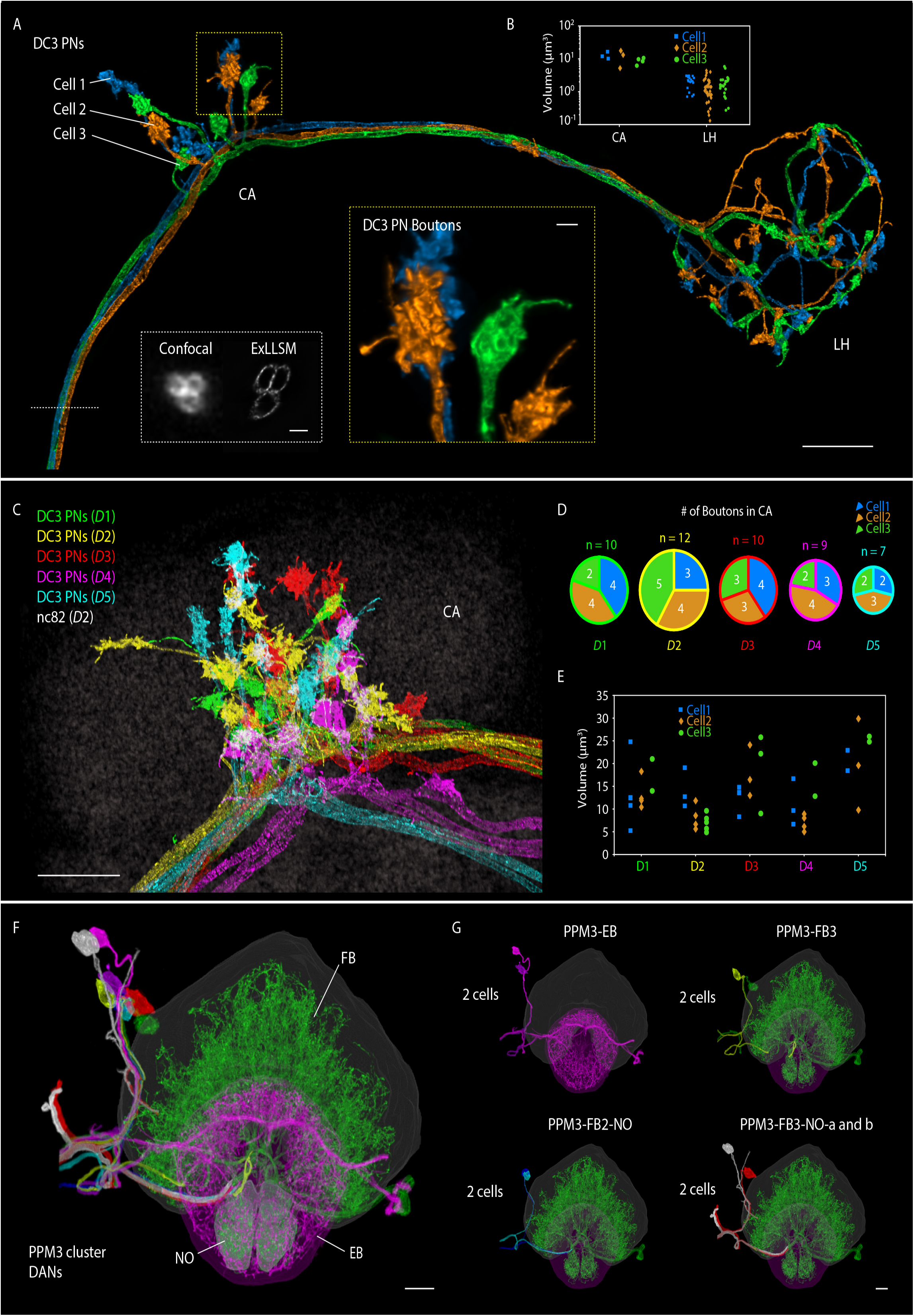
Long-range tracing and stereotypy of neuron bundles in *Drosophila*. (**A**) MIP view of DC3 olfactory projection neurons (PNs) projecting from antenna lobe of an adult fly and partially traced here (Movie 7) to the calyx (CA) and lateral horn (LH). Scale bar, 10 μm. Inset (white box) compares cross-sectional views of the axon bundle by confocal microscopy (left) and ExLLSM (right). Scale bar, 1 μm. Inset (yellow box), shows a magnified view of DC3 PN boutons in CA. Scale bar, 1 μm. (**B**) Volume of each individual DC3 PN bouton in CA and LH. (**C**) Overlaid MIP view of DC3 PNs from five adult *Drosophila* brains (*D1*-*D5*) near CA. Scale bar, 10 μm. (**D**) Number of DC3 PN boutons in CA for *D1*-*D5* shown in (C). (**E**) Volume of DC3 PN boutons in CA for *D1*-*D5* shown in (C). (**F**) MIP view of individually traced PPM3 dopaminergic neurons (DANs) in the right hemisphere of an adult *Drosophila* brain (Movie 8), innervating the fan-shaped body (FB, green), ellipsoid body (EB, magenta) and noduli (NO, green). The fine neurites arboring FB, EB and NO are from both hemispheres of the brain. Scale bar, 10 μm. (**G**) MIP view of the identified cell types of PPM3 DANs. See also fig. S18. Scale bar, 10 μm.

**Fig. 6.**
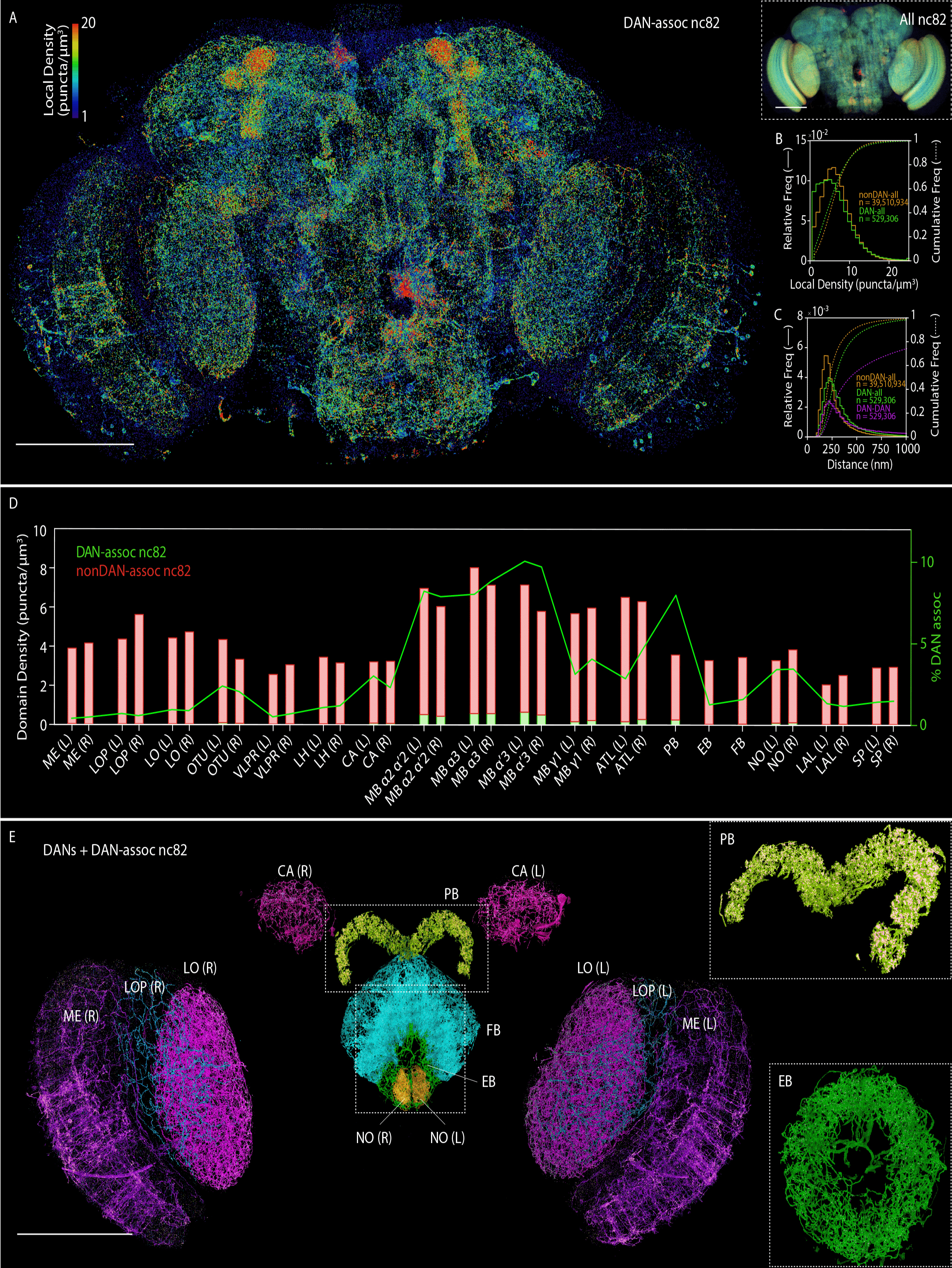
Whole brain analysis of presynaptic sites and dopaminergic neurons in *Drosophila*. (**A**) MIP view of the subset of nc82 puncta marking presynaptic sites that are associated with dopaminergic neurons (DAN-assoc nc82), color coded by the local puncta density, in an adult *Drosophila* brain (Movie 9). Scale bar, 100 μm. Inset (right), MIP view of all nc82 puncta, using identical color coding of local density. Scale bar, 100 μm. (**B**) Distribution of local densities of DAN-associated nc82 puncta (green) and nonDAN-associated nc82 puncta (orange) in (A). See also fig. S23. (**C**) Distribution of distances from DAN-associated nc82 puncta (green) and nonDAN-associated nc82 puncta (orange) to the nearest nc82 punctum of any kind, and nearest-neighbor distances from one DAN-associated nc82 to another (magenta). See also fig. S24. (**D**) Volumetric density of DAN-associated nc82 puncta (green bars) and nonDAN-associated nc82 puncta (red bars), and the percentage of nc82 puncta that are DAN-associated (green curve), within each of the 33 neuropil domains of the adult *Drosophila* brain. See also fig. S25. (**E**) MIP view of DANs and DAN-associated nc82 puncta, color-coded by 13 representative neuropil domains (Movie 10). Scale bar, 100 μm. Insets show magnified views of the PB (top, angled view) and EB (bottom). Neuropil domains: ME (medulla), LOP (lobula plate), LO (lobula), OTU (optical tubercle), VLPR (ventrolateral protocerebrum), LH (lateral horn), CA (calyx), MB (mushroom body), ATL (antler), PB (protocerebral bridge), EB (ellipsoid body), FB (fan-shaped body), NO (noduli), LAL (lateral accessory lobe), and SP (superior protocerebrum). L and R indicate the left and right hemisphere of the brain, respectively.

The neuronal circuits of the olfactory pathways to the mushroom body (MB) have been extensively described using light microscopy and have been reconstructed completely in the L1 instar larva and partially in the adult brain using EM (*9*, *75*). However, the variation among individual animals has not been well studied at the level of detailed subcellular circuitry. The speed of ExLLSM now makes this possible. We therefore studied the stereotypy of DC3 PNs by comparing their morphologies in the CA across five different animals (Fig. 5C). As expected, we consistently observed more boutons in LH than in CA, larger boutons in CA, and the restriction of boutons to the ends of the neurites in CA. However, we also found that both the number and size of boutons differ among the three cells from the same hemisphere as well as between animals. For example, the total number of boutons in CA varied from 7 to 12 and none of the bouton assignments to each cell was the same among all five brains studied (Fig. 5D). The bouton size also showed substantial variability among the brains (Fig. 5E). These variations might arise from the distinct developmental histories of the individual animals, but it is not yet clear whether they also indicate differences in synaptic strength and connection with Kenyon cells, or how they might affect processing of olfactory information for associative learning in the mushroom body. ExLLSM enables future experiments to address such questions, thanks to its high throughput and the precise descriptions of neuronal morphology that it delivers.

Given our success with this relatively simple example, we next applied ExLLSM to a much more challenging sample by imaging a ~340 × 660 × 90 μm^3^ volume covering nearly the entire brain of a TH-GAL4 transgenic *Drosophila* specimen immunostained in one color against the membranes of all dopaminergic neurons (DANs) and in a second color with nc82 antibodies against Bruchpilot (Brp), a major structural and functional component of presynaptic active zones (AZs) (*76*, *77*). Among the ~110 DANs within the image volume, we focused our efforts on tracing the protocerebral posterior medial 3 (PPM3) cluster of DANs that project to the central complex, a key brain region essential for navigation, visual memory, sleep and aggression (*78*–*80*). With manual annotation, we identified and traced all eight individual cells within the cluster (Fig. 5F, fig. S18, Movie 8). While tracing of fine processes inside the central complex was difficult, we were able to trace the main axonal branches and precisely determine the number of cell types and the number of cells belonging to each cell type. Within the PPM3 cluster, we found that two cells (PPM3-EB) mainly projected to the ellipsoid body (EB) (*80*), two cells (PPM3-FB3) projected to layer 3 of the fan-shaped body (FB), two cells (PPM3-FB2-NO) projected to layer 2 of the FB and noduli (NO), and two cells, which could be further categorized into two cell types (PPM3-FB3-NO-a and PPM3-FB3-NO-b), projected to layer 3 of the FB and NO (Fig. 5G, fig. S18, supplementary note 6f). Using stochastic labeling of individual neurons and split-GAL4 intersection, we were able to identify and confirm the individual cell types we assigned (fig. S18, supplementary note 6f). This determination will provide a basis to design and interpret future experiments.

### Whole Brain Analysis of Presynaptic Sites and Dopaminergic Neurons

We next turned our attention to the nc82 channel of this specimen, since recent EM measurements of the nearest-neighbor distances between synapses in the α lobe of the MB (fig. S19) (*81*) suggest that quantitative counting of synapses across the *Drosophila* brain should be possible with ExLLSM at 4× expansion. However, to have confidence in the results, we needed to show that nc82 puncta larger than 100 nm represented true AZs and not nonfunctional Brp monomers or non-specific background. To do so, we imaged two additional nc82 stained brains, one co-immunostained against V5-tagged Brp, and the other co-immunostained against the AZ protein Syd1 (supplementary note 6c). In both cases, the distribution of distances from each nc82 punctum to its nearest co-stained neighbor was consistent with their mutual incorporation in a single AZ (fig. S20). As additional confirmation, we returned to the DAN / nc82 specimen and measured a 70-fold higher surface density of nc82 puncta at the axons and boutons of the output neuron from the α1 compartment of the MB (MBON-α1) than at its dendrites (fig. S21, supplementary note 6d), consistent with the near-absence of dendritic presynaptic densities observed for the same neuron by EM (*81*). Furthermore, we counted ~44,000 nc82 puncta in the α3 compartment (fig. S22), compared to ~34,000 synapses in the EM study (fig. S19), with the difference possibly attributable to the different sexes and ages of the animals studied (supplementary note 6e). The distribution of synaptic distances was also similar in the two cases (figs. S19B, S22B).

Given confidence from these results, we then extended our analysis across nearly the entire brain (the medial lobes of the MB were not imaged because TH-Gal4 does not express in the DAN in that region). In total, we counted ~40 million nc82 puncta, ~530,000 of them localized at DAN (Fig. 6A, Movie 9), and calculated the brain-wide distribution of puncta density (Fig. 6B) and nearest neighbor distances between any puncta or only DAN-associated ones (Fig. 6C).

Notably, we observed substantial differences when we further subdivided our analysis into 33 major neuropil domains (figs. S23-S25, Table S5). The volume density of all puncta, for example, varied from ~2-3 per μm^3^ in the lateral accessory lobe (LAL) and superior protocerebrμm (SP) to ~6-8 in the compartments of the MB (Fig. 6D), perhaps reflecting the distinct computational needs of different brain regions. The high density in the MB, for instance, is likely beneficial for increasing capacity and sensory specificity of memory in associative learning.

When focusing on only those nc82 puncta associated with DAN, we found additional differences. For example, the distance between non-DAN nc82 puncta and DAN-associated nc82 puncta differed substantially between neuropil domains (fig. S24), indicating that proportion of synapses that can be modulated by dopamine may differ between brain regions. We also found that the percentage of puncta associated with DAN was approximately tenfold higher in the MB than in the optic lobes (Fig. 6D), consistent with observation that dopamine dependent heterosynaptic plasticity is the basis of associative learning in the MB (*81*–*83*). On the other hand, the FB and the EB, which are known for visual and place memory formation (*84*), exhibited surprisingly low DAN association, while the protocerebral bridge (PB) and the antler (ATL), which are not particularly known for heterosynaptic plasticity, showed high DAN association second only to the MB. Despite these differences, the variation in surface density of nc82 puncta on DAN in different neuropil domains was considerably less pronounced (fig. S25B), as the percentage volume occupied by DAN in each domain (fig. S25D) followed similar trends to the percentage of DAN-associated puncta (Fig. 6D). This can also be seen directly in volume renderings of the DAN and DAN-associated puncta in each neuropil domain (Fig. 6E, Movie 10), although local intra-domain variations in the spatial distribution of nc82 can also be seen.

## Discussion

Thanks to its combination of high imaging speed, low photobleaching rate, and 3D nanoscale resolution, ExLLSM extends, by at least 1000-fold in volume, the ability of SR fluorescence microscopy to generate detailed images of subcellular ultrastructure, filling a valuable niche between the high throughput of conventional optical pipelines of neural anatomy (*12*, *13*) and the ultrahigh resolution of corresponding EM pipelines (*9*, *68*, *81*). With genetically targeted cell-type specific labeling (*21*, *85*–*87*) and protein-specific immunostaining, ExLLSM enables sparse neural subsets and dense synaptic connections to be recorded, visualized, and quantified at ~60 × 60 × 90 nm^3^ resolution with ~100 person-hours of effort over cortex spanning volumes in the mouse (Figs. 3 & 4) or brain-wide volumes in *Drosophila* (Fig. 6), compared to five weeks to image and ~16,000 person-hours to trace all neurons and count all synapses in an 80-fold smaller volume encompassing the α lobe of the MB in a recent EM study at 8 nm isotropic resolution (*81*). The fluorescence contrast of ExLLSM also raises the possibility of correlating (*88*) fluorescence-based genetic indicators of neural activity (*89*, *90*) with neural ultrastructure over much larger volumes and without the labeling compromises common to correlative EM/fluorescence studies (*91*).

Although we have focused on the mouse cortex and the *Drosophila* brain in this work, we have also applied ExLLSM to image the mossy fiber innervation of granule cells in glomeruli in the cerebellum of the mouse (fig. S26, movie S3), as well as a complete human kidney glomerulus section (fig. S27). However, the application of ExM to any biological system must be examined on a case-by-case basis through careful controls and comparisons to known aspects (e.g., by EM) of the specific ultrastructural elements under investigation. In particular, extrapolating the faithful nanoscale expansion of delicate membranous structures and vesicles in a specimen from images of more robust components such as cytoskeletal elements, clathrin-coated pits, or nuclear histones (*22*, *45*, *92*, *93*) should be avoided. Elastic inhomogeneity of the specimen post-digestion such as from collagen-rich connective tissue or adhesion to a rigid substrate can also interfere with expansion. In this regard, brain tissue may represent a best case for ExM studies, due to its comparatively homogenous mechanical properties and ready digestion. It should always be remembered that any image of a once-living specimen is an imperfect representation of that specimen, and the more steps that intrude in the process from one to the other, the more imperfect it becomes. Overexpression, chemical fixation, permeabilization, and immunostaining already introduce numerous structural artifacts (*94*–*96*) in all forms of high resolution fluorescence microscopy including ExM, but ExM also requires additional steps of polymer infusion, gelation, label attachment, digestion, expansion, and handling that can perturb ultrastructure even more. Careful controls are essential.

At 4× expansion, the resolution of ExLLSM is close, but not quite sufficient, to trace fine, highly innervated neuronal processes such as the PPM3 cluster terminating in the central complex (Fig. 5F) or the dorsal paired medial (DPM) neurons that innervate the MBs (movie S4), and would therefore benefit from higher expansion ratios. However, while up to 20× expansion has been reported (*45*), our efforts to expand mouse cortical tissue to 8× (supplementary note 1e.1) has revealed regions where the expansion superficially appears accurate (fig. S28A), and other regions of clear distortion, such as irregularly shaped somata and nuclei (fig. S28B).

Even if specimen-wide isotropic expansion can be achieved at higher ratios, ExM is still heir to the problems that bedevil other forms of high resolution fluorescence microscopy. Chief among these is that, due to the stochastic nature of labeling, the mean separation between fluorophores must be ~5-10× smaller than the desired resolution in each dimension in order to distinguish with high confidence two or more structures for which no *a priori* knowledge exists (*97*). We met this requirement at the level of ~60 × 60 × 90 nm resolution in most cases thanks to the dense expression of cytosolic label in Thy1-YFP transgenic mice (Fig. 1-4) and DAN membrane label in a TH-Gal4 transgenic fly (Fig. 5, 6), as well as the exceptional specificity of ABs targeting MBP (Fig. 2, 4) and nc82 (Fig. 6). Other ABs in our study did not meet this standard, but were sufficient to identify organelles responsible for voids of cytosolic label (Tom 20 and LAMP1, Fig. 2), mark Homer1 at synapses (Fig. 3) and Caspr at nodes of Ranvier (Fig. 4), and measure statistical distributions of synapse breadth and pre- and post-synaptic separation (Homer1 and Bassoon, Fig. 4). However, immunostaining in any form is probably not dense enough to achieve true 3D resolution much beyond that already obtainable at 4× expansion, and the long distance between epitope and fluorophore, particularly with secondary ABs, further limits resolution. Likewise, loss of FP fluorescence upon linking and digestion, as well as the slow continued loss of fluorescence we observed post-expansion (supplementary notes 2c and 2d), probably preclude study at high resolution of many FP-linked proteins at the endogenous levels produced by genome editing. Indeed, even at 4× expansion, we rarely found sufficient residual fluorescence to image targets labeled with red FPs of the *Anthozoa* family, despite reports to the contrary (*23*).

Despite these challenges and limitations, the high speed and nanometric 3D resolution of ExLLSM make it an attractive tool for comparative anatomical studies, particularly in the *Drosophila* brain. For example, while we imaged the entire TH-Gal4 / nc82 brain in Fig.6 in 62.5 hours (3.2 × 10^5^ μm^3^/hr), with subsequent improvements in scanning geometry and FOV we imaged mouse brain tissue (left column, Fig. 1) in two colors at 4.0 × 10^6^ μm^3^/hr. If transferrable to the fly, this would allow whole brain imaging in ~5.0 hrs. Importantly, this limit is not fundamental – with simultaneous multicolor imaging and multiple cameras to cover even broader FOVs, rates up to ~10^8^ μm^3^/hr may be achievable, or ~12 min/fly brain at 4× expansion. Assuming the future development of: a) robust, isotropic expansion at 10× or greater; b) longer working distance high NA water immersion objectives or lossless sectioning (*98*) of expanded samples; and c) a ubiquitous, dense, and cell-permeable fluorescent membrane stain analogous to heavy metal stains in EM, even densely innervated circuits might be traced, particularly when imaged in conjunction with cell-type specific or stochastically expressed multicolor labels for error checking (*99*). With such a pipeline in place, ~10-100 specimens might be imaged in a single day at 4-10× expansion, enabling statistically rich, brain-wide studies with protein-specific contrast and nanoscale resolution of neural development, sexual dimorphism, degree of stereotypy, and structure/function or structure/behavior correlations, particularly under genetic or pharmacological perturbation.

## Materials and Methods

### Preparation of ExM samples

Mouse, *Drosophila melanogaster* and human samples were dissected, fixed and immunostained following the protocols in supplementary note 2. Sample genotypes and antibodies are summarized in Table S1. Unless otherwise noted, all samples were processed using a protein-retention ExM (proExM) protocol with minor modifications (*23*, *100*) or an expansion pathology (ExPath) protocol (*101*). Prepared ExM samples were stored in 1x PBS at 4°C and expanded in doubly deionized water immediately before imaging by LLSM.

### Lattice light sheet imaging

With the exception of Fig. 1, all ExM samples were imaged in objective scan mode (*24*) using a LLSM described previously (*102*), except with adaptive optics capability disabled. The ExM sample in the left column of Fig.1 was imaged using an LLSM optimized for ExM, featuring a broader 160 μm FOV, a 1.5 mm scan range, and software optimized for rapid sample scan acquisition (supplementary note 2a). All expanded samples were large compared to the LLS FOV, and were therefore imaged in a series of overlapping 3D tiles that covered the desired sample volume (supplementary note 2b). For imaging sessions of several hours or more, focus was maintained by periodic imaging of reference beads. Raw data from each tile was deskewed (for sample scan mode), flat-fielded, deconvolved, and stored for subsequent processing.

### Computing pipeline for flat-field correction, stitching, and export of 3D image tiles

Since automatic tools for 3D stitching (*103*–*107*) do not scale to datasets with thousands of 3D image tiles, we developed a scalable high-performance computing (HPC) pipeline to robustly flat-field correct, deconvolve, and assemble 3D image tiles into the final volume (supplementary note 3). First, we extended and parallelized CIDRE (*103*) for 3D volumes to calculate 3D flat-fields (Fig. S2 and S3). We then corrected the raw image tiles using these flat-fields and deconvolved each. Next, we parallelized the globally optimizing 3D stitching method (*104*) to automatically stitch the thousands of raw image tiles, without manual intervention, in an iteratively refined prediction model that corrects for systematic stage coordinate errors (Fig. S4). Lastly, we exported the stitched datasets using the flat-field corrected and deconvolved image tiles as multi-resolution hierarchies into a custom file format (N5) (*25*) that enabled parallel block-wise export and compression on a HPC cluster. Bindings for N5 format for the ImgLib2 library (*108*) are provided for the ImageJ distribution Fiji (*109*). For interactive visualization, we developed a BigDataViewer (*110*) based viewer plugin including a crop and export tool to make arbitrary sub-volumes available in legacy formats such as TIFF image series.

## Movie Captions

**Movie 1.**
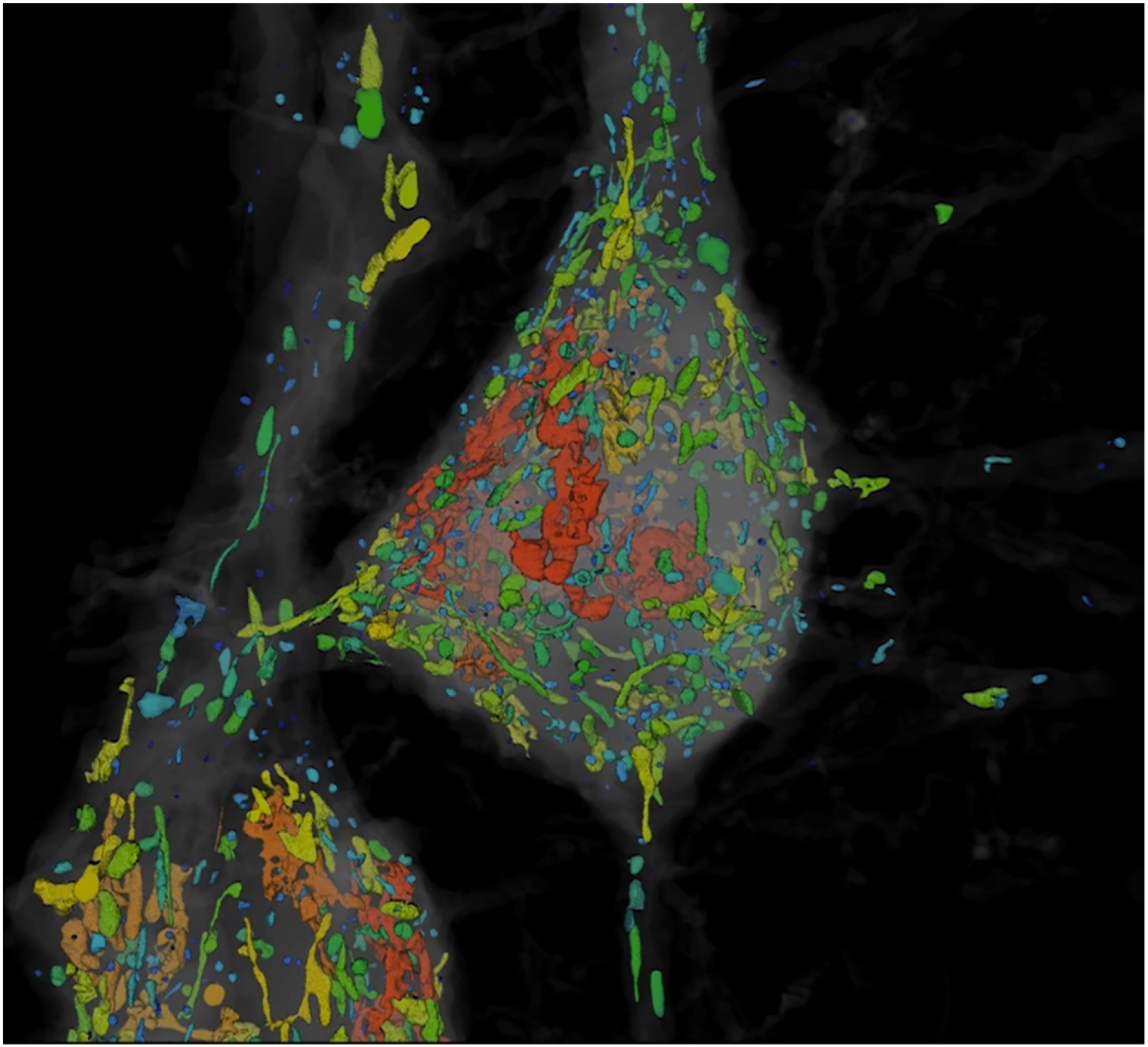
Organelle analysis in layer V pyramidal neurons in S1. Segmentation of cytosolic voids in Thy1-YFP expressing neurons, quantification of their volumes, and immunostaining-based classification of those voids that represent mitochondria or multivesicular bodies / autolysosomes (Fig. 2A-E, fig. S7, movie S1).

**Movie 2.**
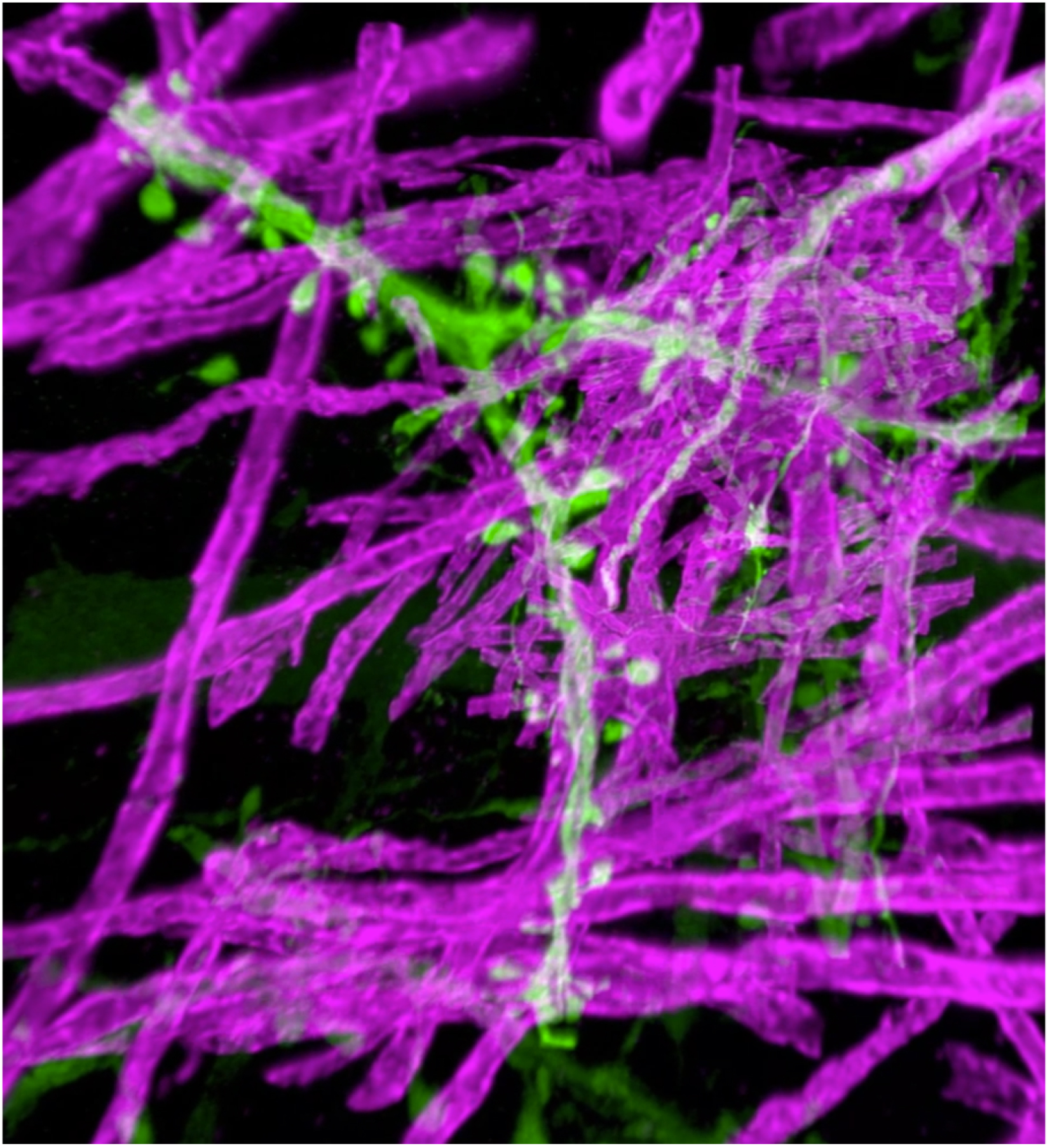
Axon myelination and local g-ratio calculation in layer V pyramidal neurons in S1. Thy1-YFP expressing neurons and immunostained myelin sheaths across 320 × 280 × 60 μm^3^, with quantification of the local g-ratio on the surface of a specific myelin sheath (Fig. 2F, G, figs. S8, S9).

**Movie 3.**
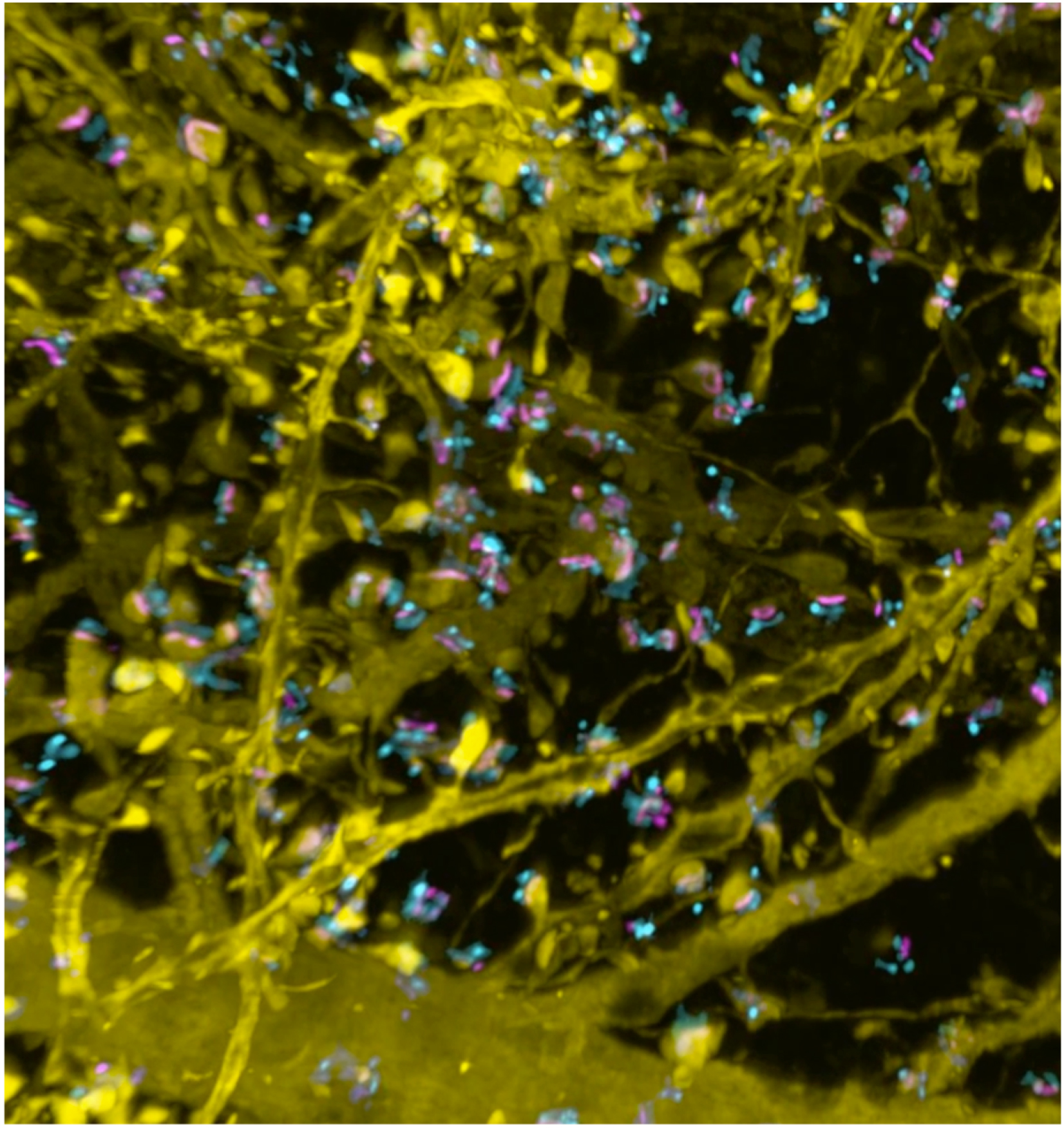
Synaptic proteins and their associations to neural processes at layers IV-V in S1. Thy1-YFP expressing neurons and immunostained pre- and post-synaptic proteins Bassoon and Homer1 across 75 × 100 × 125 μm^3^, sequentially showing all Bassoon and Homer1 puncta, and only YFP-associated Bassoon and Homer1 pairs (Figs. 2I-K, fig. S10).

**Movie 4.**
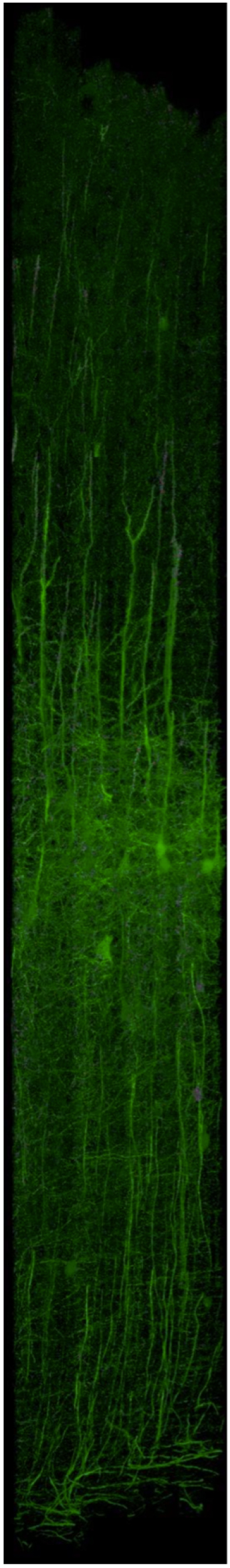
Relationship of post-synaptic Homer1 to neural processes across S1. Thy1-YFP expressing neurons and immunostained post-synaptic protein Homer1 across 1900 × 280 × 70 μm^3^, in S1 with specific focus on two adjacent layer V pyramidal neurons exhibiting substantially different patterns of Homer1 expression (Fig. 3, fig. S11-14, movie S2).

**Movie 5.**
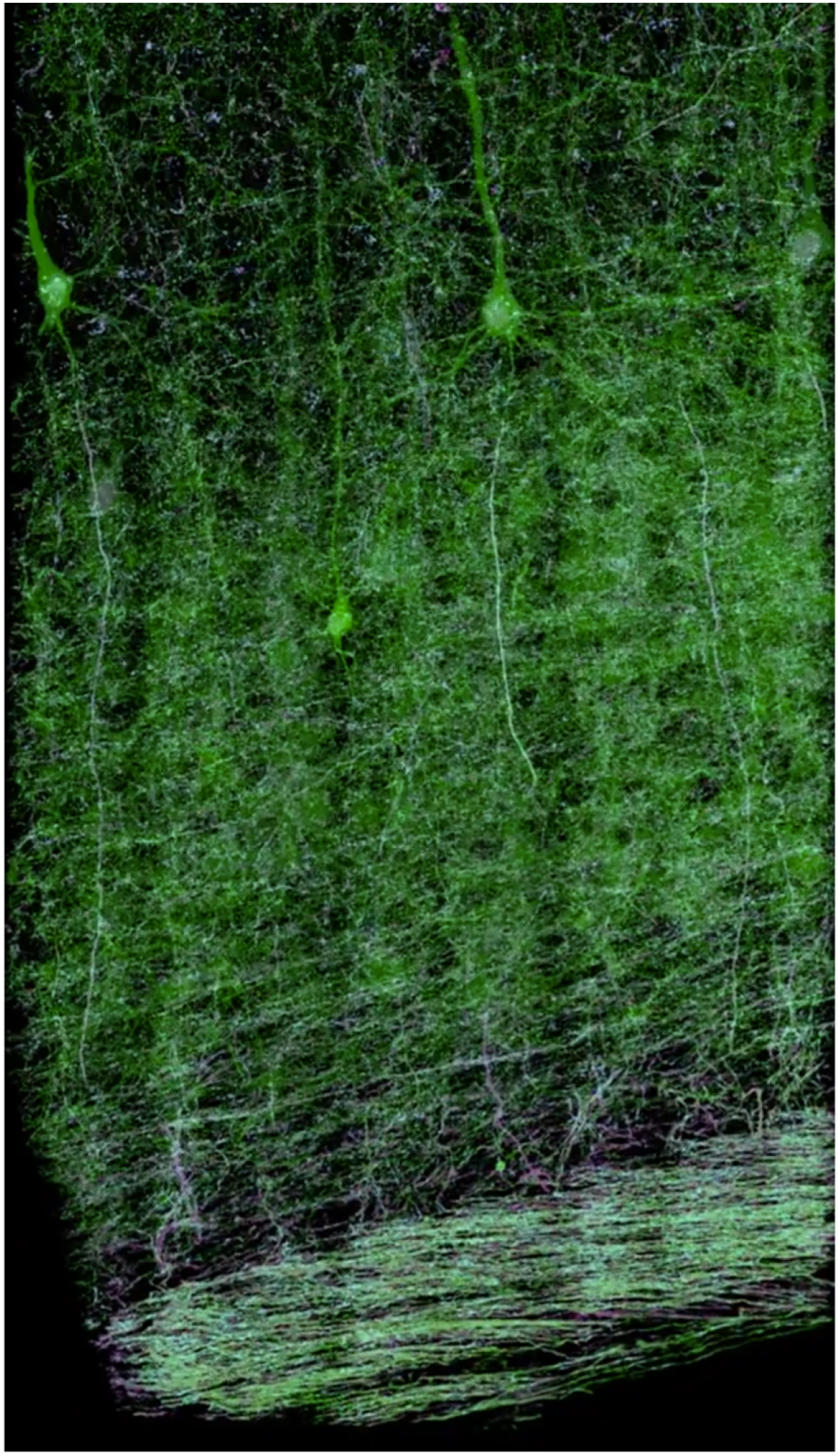
Neural processes and myelination patterns across V1. Thy1-YFP expressing neurons across 1100 × 280 × 83 μm^3^ in V1, immunostained against myelin and Caspr, a marker of the nodes of Ranvier, with specific emphasis on the neural processes and longitudinal myelination profile of a selected layer V pyramidal neuron (Fig. 4, figs. S16, S17).

**Movie 6.**
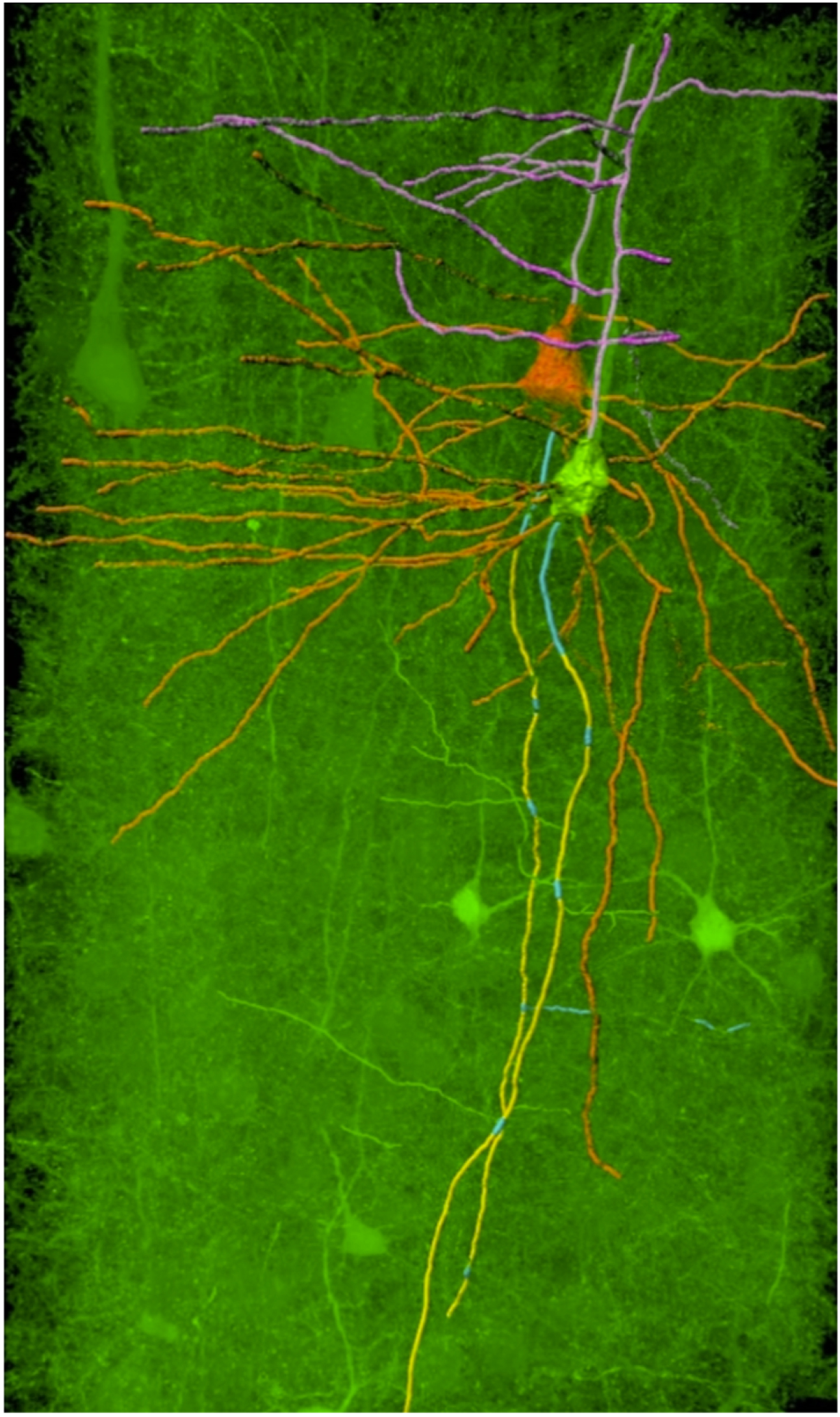
Segmentation of pyramidal neurons in layer V of V1. Segmentation of two neurons, with specific emphasis on their branches and axonal myelination patterns (Fig. 4, figs. S16, S17).

**Movie 7.**
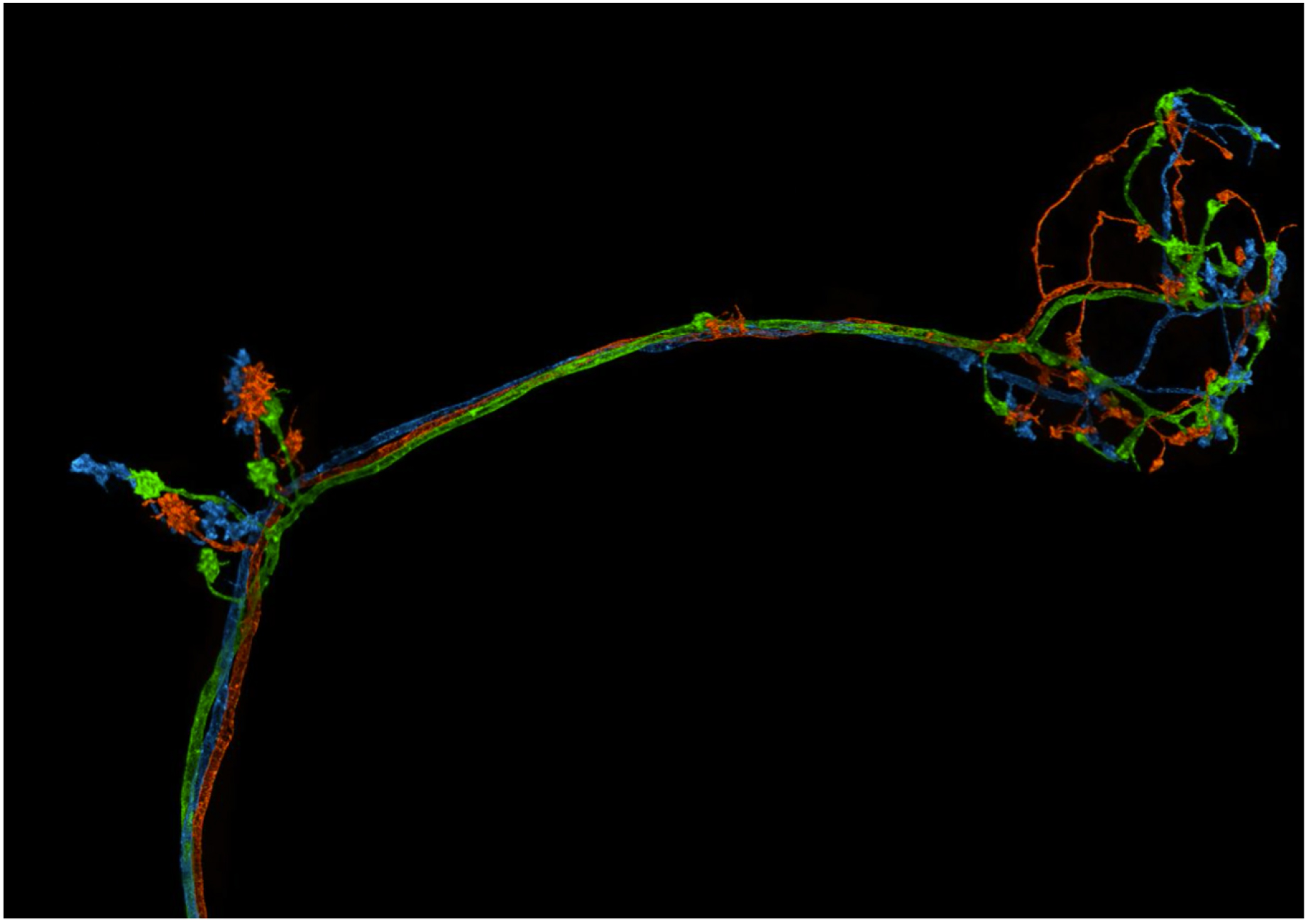
Tracing of DC3 olfactory projection neurons (PNs) in an adult *Drosophila* brain. Volumetric view of three individually traced neurons projecting from the antenna lobe in a bundle, with magnified views of their boutons at the calyx and lateral horn (Fig. 5A-E).

**Movie 8.**
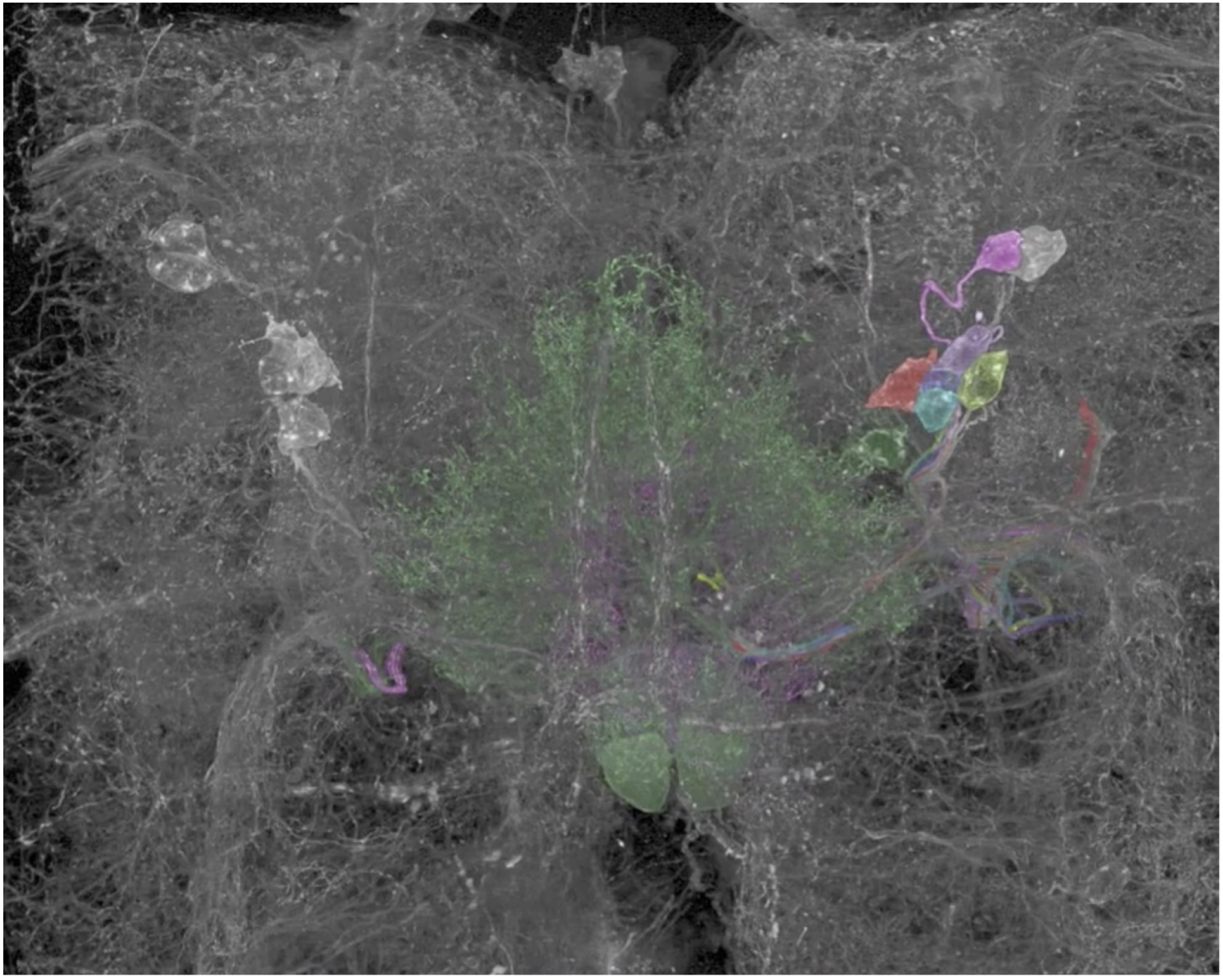
Tracing and classification of PPM3 dopaminergic neurons (DANs) in an adult *Drosophila* brain. Section of brain near the central complex with eight neurons from the protocerebral posterior medial 3 cluster in the right hemisphere (colored) shown in relation to surrounding DANs (white), and tracing of the individual neurons to their paired innervations in different regions of the central complex (Fig. 5F, G, fig. S18).

**Movie 9.**
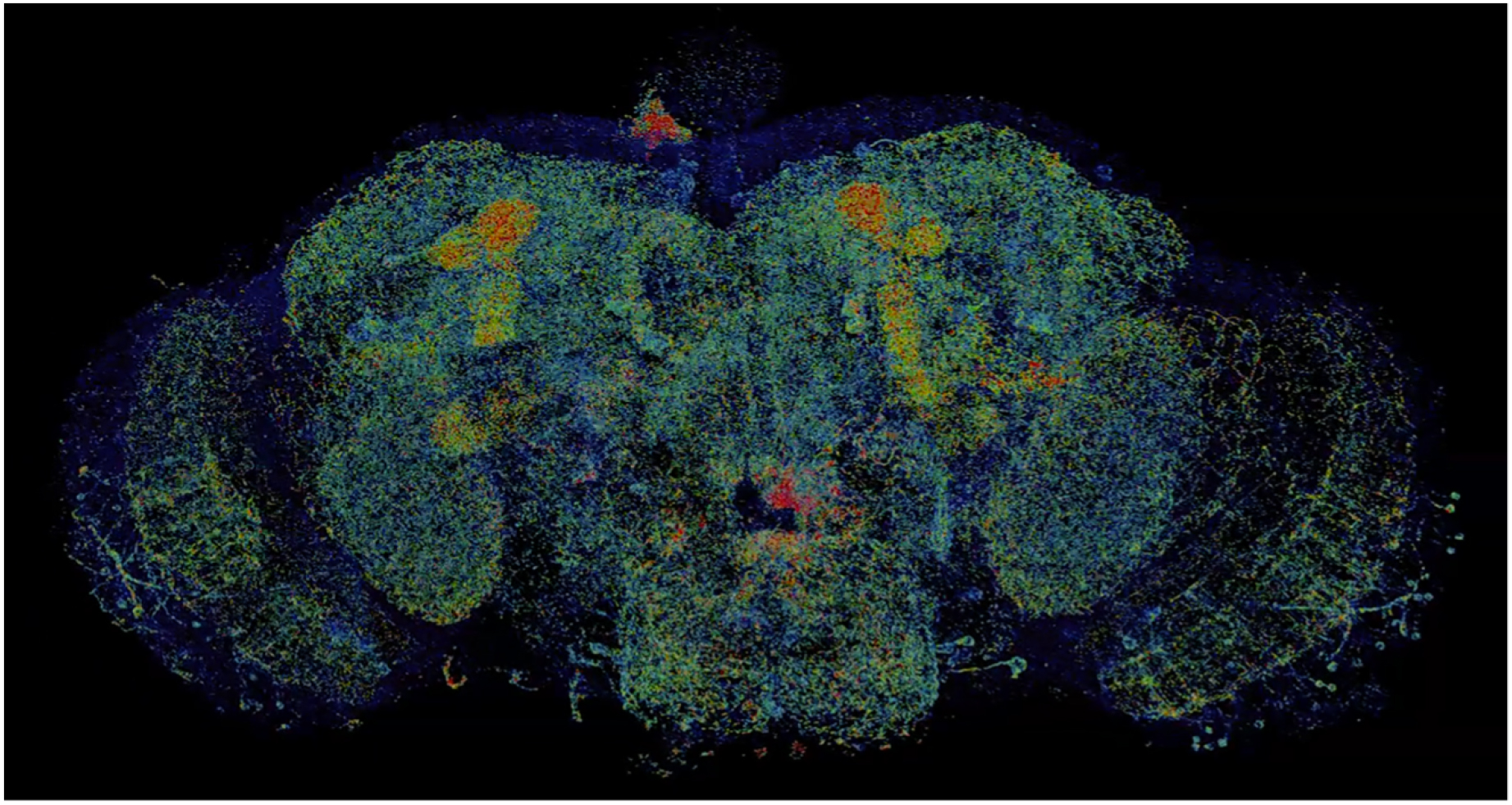
Local density map of DAN-associated presynaptic sites across an adult *Drosophila* brain. Color-coded neuropil domains and 3D color-coded map of the local density of DAN-associated nc82 puncta in each domain (Fig. 6A-D, fig. S23-25).

**Movie 10.**
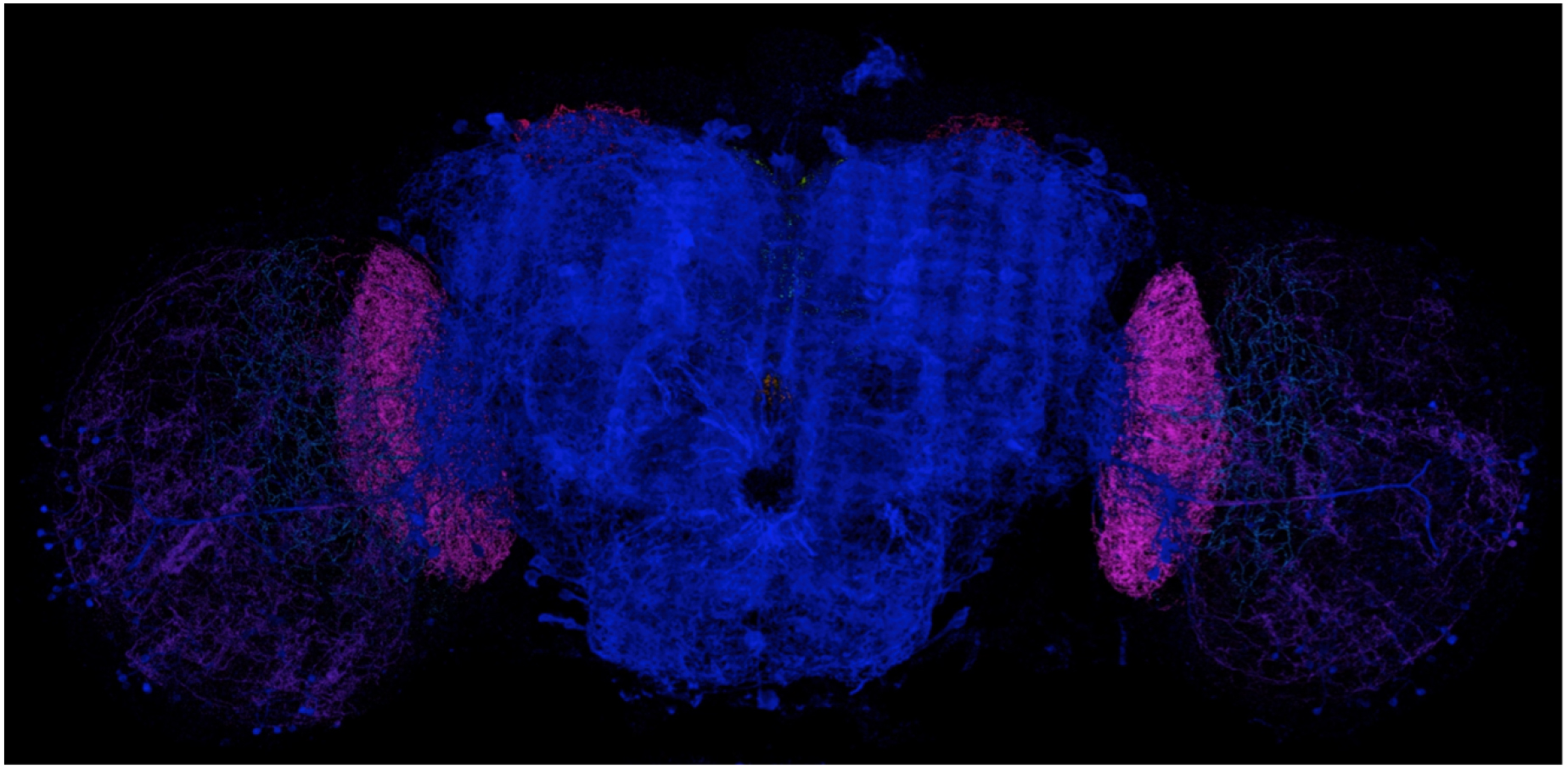
DANs and DAN-associated presynaptic sites in different neuropil domains of an adult *Drosophila* brain. Volume rendering DANs, DAN-associated nc82 puncta, and all nc82 puncta across the entire brain, color coded by neuropil domain, followed by magnified 3D and orthoslice views of DANs and DAN-associated nc82 in each of nine different domains (Fig. 6E).

## Acknowledgments

We thank D. Bock, K. Svoboda, N. Ji, N. Spruston, L. Scheffer, E. Snapp, P. Tillberg, L. Lavis, E. Bloss, W. Legant, D. Hoffman, K. Hayworth and H. Hess at Howard Hughes Medical Institute (HHMI) Janelia Research Campus (JRC) and B. Sabatini at Harvard Medical School (HMS) for invaluable discussions and comments. We also thank K. Schaefer, T. Wolff, C-L. Chang and H. Choi at JRC for help with sample preparation and imaging. We gratefully acknowledge the shared resources and project teams at JRC, including D. Alcor, J. Heddleston, and A. Taylor of the Advanced Imaging Center and Light Microscopy Facility for help with imaging, I. Negrashov and jET for manufacturing expertise, O. Malkesman, K. Salvesen, C. Christoforou, G. Meissner, and the FlyLight project team for sample handling and preparation. Lastly, we are grateful to C. Pama and R. Karadottir at University of Cambridge, J. Melander and H. Zhong at OHSU, E. Karagiannis, J-S. Kang, and F. Chen at MIT for help with sample preparation, and H. Otsuna, T. Kawase and E. Bas at JRC, C. Wietholt at FEI Amira, M. Gastinger at Bitplane Inc., J. McMullen and T. Tetreault at MBF Bioscience for data analysis and visualization.

## Author contributions

E.B., E.S.B., and R.G. supervised the project and wrote the manuscript with input from all coauthors. T.-L.L. and J.C. built the microscopes with input from E.B., D.E.M., and jET (JRC) and performed all microscope characterization experiments. D.E.M. created the instrument control software. R.G., S.M.A., T.-L.L., V.S., J.C., and C.O. acquired all biological data with coauthors. G.H.H. provided the Thy1-YFP mice and R.G. and S.M.A. prepared the ExM samples. A.G. and A.H. provided the Slc17a7-cre X TCGO mice and prepared the ExM samples. Y.Z. provided the human kidney sections and prepared the ExPath samples. Y.A. and G.M.R. created the split-GAL4 fly strains, Y.A. optimized the IHC conditions, and R.G. and S.M.A prepared the ExM samples. K.R.M., S.G.M., S.M.A., and T.-L.L. provided the initial stitching software packages. I.P. and S.S. created the automated flat-field, stitching, and N5 visualization pipeline and I.P., R.G., J.B., and S.S. deconvolved, flat-fielded, and stitched all image data. Y.A. performed segmentation and tracing, and supervised analyses of all fly image data. C.Z., S.T., and A.R. provided customized commercial software packages and helped with segmentation, tracing and reconstruction of neurites and dendritic spines using these packages. S.U. and R.G. processed and performed quantitative analysis of all image data. S.U., R.G., E.B., and Y.A. produced all figures and movies.

## Data and materials availability

All data needed to evaluate the conclusions in the paper are present in the paper or the supplementary materials. Documentation for construction of a lattice light-sheet microscope can be obtained by execution of a research license agreement with HHMI.

## References and Notes

1. T. C. Licquia, J. A. Newman, Improved estimates of the Milky Way’s stellar mass and star formation rate from hierarchical Bayesian meta-analysis. Astrophys. J. 806, 96 (2015).

2. C. J. Conselice, A. Wilkinson, K. Duncan, A. Mortlock, The evolution of galaxy number density at z < 8 and its implications. Astrophys. J. 830, 83 (2016).

3. S. Herculano-Houzel, The human brain in numbers: A linearly scaled-up primate brain. Front. Hum. Neurosci. 3, 31 (2009).

4. K. Sharma et al., Cell type- and brain region-resolved mouse brain proteome. Nat. Neurosci. 18, 1819–1831 (2015).

5. R. Yuste, The discovery of dendritic spines by Cajal. Front. Neuroanat. 9, 18 (2015).

6. J. W. Lichtman, W. Denk, The big and the small: Challenges of imaging the brain’s circuits. Science. 334, 618–623 (2011).

7. J. E. Heuser, T. S. Reese, Evidence for recycling of synaptic vesicle membrane during transmitter release at the frog neuromuscular junction. J. Cell Biol. 57, 315–344 (1973).

8. C. S. Xu et al., Enhanced FIB-SEM systems for large-volume 3D imaging. Elife. 6, e25916 (2017).

9. Z. Zheng et al., A complete electron microscopy volume of the brain of adult Drosophila melanogaster. bioRxiv (2017), doi:https://doi.org/10.1101/140905.

10. K. D. Micheva, S. J. Smith, Array tomography: A new tool for imaging the molecular architecture and ultrastructure of neural circuits. Neuron. 55, 25–36 (2007).

11. J.-C. Rah et al., Thalamocortical input onto layer 5 pyramidal neurons measured using quantitative large-scale array tomography. Front. Neural Circuits. 7, 177 (2013).

12. A. S. Chiang et al., Three-dimensional reconstruction of brain-wide wiring networks in Drosophila at single-cell resolution. Curr. Biol. 21, 1–11 (2011).

13. A. Jenett et al., A GAL4-driver line resource for Drosophila neurobiology. Cell Rep. 2, 991–1001 (2012).

14. M. N. Economo et al., A platform for brain-wide imaging and reconstruction of individual neurons. Elife. 5, e10566 (2016).

15. S. Shah, E. Lubeck, W. Zhou, L. Cai, In situ transcription profiling of single cells reveals spatial organization of cells in the mouse hippocampus. Neuron. 92, 342–357 (2016).

16. J. R. Moffitt et al., High-throughput single-cell gene-expression profiling with multiplexed error-robust fluorescence in situ hybridization. Proc. Natl. Acad. Sci. 113, 11046–11051 (2016).

17. J. Tønnesen, U. V. Nägerl, Superresolution imaging for neuroscience. Exp. Neurol. 242, 33–40 (2013).

18. H. Zhong, Applying superresolution localization-based microscopy to neurons. Synapse. 69, 283–294 (2015).

19. C. I. Bargmann, Beyond the connectome: How neuromodulators shape neural circuits. BioEssays. 34, 458–465 (2012).

20. J. Lu, J. C. Tapia, O. L. White, J. W. Lichtman, The interscutularis muscle connectome. PLoS Biol. 7, 0265–0277 (2009).

21. A. Nern, B. D. Pfeiffer, G. M. Rubin, Optimized tools for multicolor stochastic labeling reveal diverse stereotyped cell arrangements in the fly visual system. Proc. Natl. Acad. Sci. 112, E2967–E2976 (2015).

22. F. Chen, P. W. Tillberg, E. S. Boyden, Expansion microscopy. Science. 347, 543–548 (2015).

23. P. W. Tillberg et al., Protein-retention expansion microscopy of cells and tissues labeled using standard fluorescent proteins and antibodies. Nat. Biotechnol. 34, 987–992 (2016).

24. B. C. Chen et al., Lattice light-sheet microscopy: Imaging molecules to embryos at high spatiotemporal resolution. Science. 346, 1257998 (2014).

25. I. Pisarev, S. Saalfeld, Stitcher and N5 viewer, (available at https://github.com/saalfeldlab/stitching-spark and https://github.com/saalfeldlab/n5-viewer).

26. J. Tønnesen, V. V. G. K. Inavalli, U. V. Nägerl, Super-resolution imaging of the extracellular space in living brain tissue. Cell. 172, 1108–1121 (2018).

27. C. J. L. Sheppard, Super resolution in confocal imaging. Optik (Stuttg). 80, 53–54 (1988).

28. C. B. Müller, J. Enderlein, Image scanning microscopy. Phys. Rev. Lett. 104, 198101 (2010).

29. X.-T. Cheng et al., Characterization of LAMP1-labeled nondegradative lysosomal and endocytic compartments in neurons. J. Cell Biol. (2018), doi:10.1083/jcb.201711083.

30. N. Kasthuri et al., Saturated reconstruction of a volume of neocortex. Cell. 162, 648–661 (2015).

31. Q. A. Liu, H. Shio, Mitochondrial morphogenesis, dendrite development, and synapse formation in cerebellum require both Bcl-w and the glutamate receptor δ2. PLoS Genet. 4, e1000097 (2008).

32. W. Kuehnel, Color Atlas of Cytology, Histology, and Microscopic Anatomy (Thieme Flexibook, ed. 4th, 2003).

33. V. Popov, N. I. Medvedev, H. A. Davies, M. G. Stewart, Mitochondria form a filamentous reticular network in hippocampal dendrites but are present as discrete bodies in axons: A three-dimensional ultrastructural study. J. Comp. Neurol. 492, 50–65 (2005).

34. M. R. Duchen, Mitochondria in health and disease: Perspectives on a new mitochondrial biology. Mol. Aspects Med. 25, 365–451 (2004).

35. S. G. Waxman, M. V. I. Bennett, Relative conduction velocities of small myelinated and non-myelinated fibres in the central nervous system. Nat. New Biol. 238, 217–219 (1972).

36. K. A. Nave, Myelination and support of axonal integrity by glia. Nature. 468 (2010), pp. 244–252.

37. J. J. Harris, D. Attwell, The energetics of CNS white matter. J. Neurosci. 32, 356–371 (2012).

38. A. Compston, A. Coles, Multiple sclerosis. Lancet. 372, 1502–1517 (2008).

39. W. A. H. Rushton, A theory of the effects of fibre size in medullated nerve. J. Physiol. 115, 101–122 (1951).

40. R. La Marca et al., TACE (ADAM17) inhibits Schwann cell myelination. Nat. Neurosci. 14, 857–865 (2011).

41. L.-J. Oluich et al., Targeted ablation of oligodendrocytes induces axonal pathology independent of overt demyelination. J. Neurosci. 32, 8317–8330 (2012).

42. M. Zonouzi et al., GABAergic regulation of cerebellar NG2 cell development is altered in perinatal white matter injury. Nat. Neurosci. 18, 674–682 (2015).

43. O. O. Glebov, S. Cox, L. Humphreys, J. Burrone, Neuronal activity controls transsynaptic geometry. Sci. Rep. 6, 22703 (2016).

44. A. Dani, B. Huang, J. Bergan, C. Dulac, X. Zhuang, Superresolution imaging of chemical synapses in the brain. Neuron. 68, 843–856 (2010).

45. J. B. Chang et al., Iterative expansion microscopy. Nat. Methods. 14, 593–599 (2017).

46. N. L. Rochefort, A. Konnerth, Dendritic spines: From structure to in vivo function. EMBO Rep. 13, 699–708 (2012).

47. H. Hering, M. Sheng, Dentritic spines: Structure, dynamics and regulation. Nat. Rev. Neurosci. 2, 880–888 (2001).

48. Esther A. Nimchinsky, Bernardo L. Sabatini, and K. Svoboda, Structure and function of dendritic spines. Annu. Rev. Physiol. 64, 313–353 (2002).

49. M. Segal, Dendritic spines and long-term plasticity. Nat. Rev. Neurosci. 6, 277–284 (2005).

50. S. Konur, D. Rabinowitz, V. L. Fenstermaker, R. Yuste, Systematic regulation of spine sizes and densities in pyramidal neurons. J. Neurobiol. 56, 95–112 (2003).

51. D. Dumitriu, A. Rodriguez, J. H. Morrison, High-throughput, detailed, cell-specific neuroanatomy of dendritic spines using microinjection and confocal microscopy. Nat. Protoc. 6, 1391–1411 (2011).

52. M. Jiang et al., Dendritic arborization and spine dynamics are abnormal in the mouse model of MECP2 duplication syndrome. J. Neurosci. 33, 19518–19533 (2013).

53. X. Yu, Y. Zuo, Two-photon in vivo imaging of dendritic spines in the mouse cortex using a thinned-skull preparation. J. Vis. Exp. 87, e51520 (2014).

54. J. I. Arellano, R. Benavides-Piccione, J. DeFelipe, R. Yuste, Ultrastructure of dendritic spines: Correlation between synaptic and spine morphologies. Front. Neurosci. 1, 131–143 (2007).

55. C. Bosch et al., FIB/SEM technology and high-throughput 3D reconstruction of dendritic spines and synapses in GFP-labeled adult-generated neurons. Front Neuroanat. 9, 60 (2015).

56. K. Takasaki, B. L. Sabatini, Super-resolution 2-photon microscopy reveals that the morphology of each dendritic spine correlates with diffusive but not synaptic properties. Front. Neuroanat. 8, 29 (2014).

57. J. Tønnesen, G. Katona, B. Rózsa, U. V. Nägerl, Spine neck plasticity regulates compartmentalization of synapses. Nat. Neurosci. 17, 678–685 (2014).

58. D. L. Dickstein et al., Automatic dendritic spine quantification from confocal data with neurolucida 360. Curr. Protoc. Neurosci. 77 (2016), doi:10.1002/cpns.16.

59. E. G. Jones, T. P. S. Powell, Morphological variations in the dendritic spines of the neocortex. J. Cell Sci. 5, 509–529 (1969).

60. J. Grutzendler, N. Kasthuri, W. B. Gan, Long-term dendritic spine stability in the adult cortex. Nature. 420, 812–816 (2002).

61. L. Anton-Sanchez et al., Three-dimensional distribution of cortical synapses: A replicated point pattern-based analysis. Front. Neuroanat. 8, 85 (2014).

62. J. DeFelipe, L. Alonso-Nanclares, J. I. Arellano, Microstructure of the neocortex: Comparative aspects. J. Neurocytol. 31, 299–316 (2002).

63. C. Sala et al., Regulation of dendritic spine morphology and synaptic function by Shank and Homer. Neuron. 31, 115–130 (2001).

64. U. Thomas, Modulation of synaptic signalling complexes by Homer proteins. J. Neurochem. 81, 407–413 (2002).

65. A. Dosemeci, R. J. Weinberg, T. S. Reese, J.-H. Tao-Cheng, The postsynaptic density: There is more than meets the eye. Front. Synaptic Neurosci. 8, 23 (2016).

66. G. H. Diering et al., Homer1a drives homeostatic scaling-down of excitatory synapses during sleep. Science. 355, 511–515 (2017).

67. D. Debanne, E. Campanac, A. Bialowas, E. Carlier, G. Alcaraz, Axon physiology. Physiol. Rev. 91, 555–602 (2011).

68. G. S. Tomassy et al., Distinct profiles of myelin distribution along single axons of pyramidal neurons in the neocortex. Science. 344, 319–324 (2014).

69. S. Einheber et al., The axonal membrane protein Caspr, a homologue of neurexin IV, is a component of the septate-like paranodal junctions that assemble during myelination. J. Cell Biol. 139, 1495–1506 (1997).

70. C. Porrero, P. Rubio-Garrido, C. Avendaño, F. Clascá, Mapping of fluorescent protein-expressing neurons and axon pathways in adult and developing Thy1-eYFP-H transgenic mice. Brain Res. 1345, 59–72 (2010).

71. S. Ramaswamy, H. Markram, Anatomy and physiology of the thick-tufted layer 5 pyramidal neuron. Front. Cell. Neurosci. 9, 233 (2015).

72. L. M. Palmer, G. J. Stuart, Site of action potential initiation in layer 5 pyramidal neurons. J. Neurosci. 26, 1854–1863 (2006).

73. S. J. C. Caron, V. Ruta, L. F. Abbott, R. Axel, Random convergence of olfactory inputs in the Drosophila mushroom body. Nature. 497, 113–117 (2013).

74. N. J. Butcher, A. B. Friedrich, Z. Lu, H. Tanimoto, I. A. Meinertzhagen, Different classes of input and output neurons reveal new features in microglomeruli of the adult Drosophila mushroom body calyx. J. Comp. Neurol. 520, 2185–2201 (2012).

75. K. Eichler et al., The complete connectome of a learning and memory centre in an insect brain. Nature. 548, 175–182 (2017).

76. W. Fouquet et al., Maturation of active zone assembly by Drosophila Bruchpilot. J. Cell Biol. 186, 129–145 (2009).

77. N. Ehmann et al., Quantitative super-resolution imaging of Bruchpilot distinguishes active zone states. Nat. Commun. 5, 4650 (2014).

78. Z. Mao, R. L. Davis, Eight different types of dopaminergic neurons innervate the Drosophila mushroom body neuropil: Anatomical and physiological heterogeneity. Front. Neural Circuits. 3, 5 (2009).

79. E. C. Kong et al., A pair of dopamine neurons target the D1-like dopamine receptor DopR in the central complex to promote ethanol-stimulated locomotion in Drosophila. PLoS One. 5, e9954 (2010).

80. O. V. Alekseyenko et al., Single serotonergic neurons that modulate aggression in Drosophila. Curr. Biol. 24, 2700–2707 (2014).

81. S. Takemura et al., A connectome of a learning and memory center in the adult Drosophila brain. Elife. 6, e26975 (2017).

82. Y. Aso et al., The neuronal architecture of the mushroom body provides a logic for associative learning. Elife. 3, e04577 (2014).

83. Y. Aso, G. M. Rubin, Dopaminergic neurons write and update memories with cell-type-specific rules. Elife. 5, e16135 (2016).

84. L. Kahsai, T. Zars, Learning and memory in Drosophila: Behavior, genetics, and neural systems. Int. Rev. Neurobiol. 99, 139–167 (2011).

85. H. Luan, N. C. Peabody, C. R. Vinson, B. H. White, Refined spatial manipulation of neuronal function by combinatorial restriction of transgene expression. Neuron. 52, 425–436 (2006).

86. B. D. Pfeiffer et al., Refinement of tools for targeted gene expression in Drosophila. Genetics. 186, 735–755 (2010).

87. M. J. Dolan et al., Facilitating neuron-specific genetic manipulations in Drosophila melanogaster using a split GAL4 repressor. Genetics. 206, 775–784 (2017).

88. D. D. Bock et al., Network anatomy and in vivo physiology of visual cortical neurons. Nature. 471, 177–184 (2011).

89. T.-W. Chen et al., Ultrasensitive fluorescent proteins for imaging neuronal activity. Nature. 499, 295–300 (2013).

90. B. F. Fosque et al., Labeling of active neural circuits in vivo with designed calcium integrators. Science. 347, 755–760 (2015).

91. P. De Boer, J. P. Hoogenboom, B. N. G. Giepmans, Correlated light and electron microscopy: Ultrastructure lights up! Nat. Methods. 12, 503–513 (2015).

92. T. J. Chozinski et al., Expansion microscopy with conventional antibodies and fluorescent proteins. Nat. Methods. 13, 485–488 (2016).

93. T. Ku et al., Multiplexed and scalable super-resolution imaging of three-dimensional protein localization in size-adjustable tissues. Nat. Biotechnol. 34, 973–981 (2016).

94. U. Schnell, F. Dijk, K. A. Sjollema, B. N. G. Giepmans, Immunolabeling artifacts and the need for live-cell imaging. Nat. Methods. 9, 152–158 (2012).

95. D. R. Whelan, T. D. M. Bell, Image artifacts in single molecule localization microscopy: Why optimization of sample preparation protocols matters. Sci. Rep. 5, 7924 (2015).

96. D. Li et al., Extended-resolution structured illumination imaging of endocytic and cytoskeletal dynamics. Science. 349, aab3500 (2015).

97. W. R. Legant et al., High-density three-dimensional localization microscopy across large volumes. Nat. Methods. 13, 359–365 (2016).

98. K. J. Hayworth et al., Ultrastructurally smooth thick partitioning and volume stitching for large-scale connectomics. Nat. Methods. 12, 319–322 (2015).

99. Y.-G. Yoon et al., Feasibility of 3D reconstruction of neural morphology using expansion microscopy and barcode-guided agglomeration. Front. Comput. Neurosci. 11, 97 (2017).

100. S. M. Asano et al., Expansion microscopy: Protocols for imaging proteins and RNA in cells and tissues. Curr. Protoc. Cell Biol. (2018), doi:10.1002/cpcb.56.

101. Y. Zhao et al., Nanoscale imaging of clinical specimens using pathology-optimized expansion microscopy. Nat. Biotechnol. 35, 757–764 (2017).

102. T. L. Liu et al., Observing the cell in its native state: Imaging subcellular dynamics in multicellular organisms. Science. 360, eaaq1392 (2018).

103. K. Smith et al., CIDRE: An illumination-correction method for optical microscopy. Nat. Methods. 12, 404–406 (2015).

104. S. Preibisch, S. Saalfeld, P. Tomancak, Globally optimal stitching of tiled 3D microscopic image acquisitions. Bioinformatics. 25, 1463–1465 (2009).

105. D. Hörl et al., BigStitcher: Reconstructing high-resolution image datasets of cleared and expanded samples. bioRxiv (2018), doi:https://doi.org/10.1101/343954.

106. M. Emmenlauer et al., XuvTools: Free, fast and reliable stitching of large 3D datasets. J. Microsc. 233, 42–60 (2009).

107. A. Bria, G. Iannello, TeraStitcher - A tool for fast automatic 3D-stitching of teravoxel-sized microscopy images. BMC Bioinformatics. 13, 316 (2012).

108. T. Pietzsch, S. Preibisch, P. Tomančák, S. Saalfeld, ImgLib2—generic image processing in Java. Bioinformatics. 28, 3009–3011 (2012).

109. J. Schindelin et al., Fiji: An open-source platform for biological-image analysis. Nat. Methods. 9, 676–682 (2012).

110. T. Pietzsch, S. Saalfeld, S. Preibisch, P. Tomancak, BigDataViewer: Visualization and processing for large image data sets. Nat. Methods. 12, 481–483 (2015).

111. I. R. Wickersham et al., Monosynaptic Restriction of Transsynaptic Tracing from Single, Genetically Targeted Neurons. Neuron. 53, 639–647 (2007).

112. H. Dionne, K. L. Hibbard, A. Cavallaro, J. C. Kao, G. M. Rubin, Genetic reagents for making split-GAL4 lines in Drosophila. Genetics. 209, 31–35 (2018).

113. L. Tirian, B. Dickson, The VT GAL4, LexA, and split-GAL4 driver line collections for targeted expression in the Drosophila nervous system. bioRxiv (2017), doi:https://doi.org/10.1101/198648.

114. S. Truckenbrodt et al., X10 Expansion Microscopy enables 25 nm resolution on conventional microscopes. EMBO Rep., e45836 (2018).

115. J. A. Bogovic, S. Saalfeld, BigWarp, (available at https://github.com/saalfeldlab/bigwarp).

116. J. A. Bogovic, P. Hanslovsky, A. Wong, S. Saalfeld, in Proceedings - International Symposium on Biomedical Imaging (2016), pp. 1123–1126.

117. F. Aguet et al., Membrane dynamics of dividing cells imaged by lattice light-sheet microscopy. Mol. Biol. Cell. 27, 3418–3435 (2016).

118. P. A. Yushkevich et al., User-guided 3D active contour segmentation of anatomical structures: Significantly improved efficiency and reliability. Neuroimage. 31, 1116–1128 (2006).

119. A. Rodriguez, D. B. Ehlenberger, P. R. Hof, S. L. Wearne, Rayburst sampling, an algorithm for automated three-dimensional shape analysis from laser scanning microscopy images. Nat. Protoc. 1, 2152–2161 (2006).

120. A. Rodriguez, D. B. Ehlenberger, D. L. Dickstein, P. R. Hof, S. L. Wearne, Automated three-dimensional detection and shape classification of dendritic spines from fluorescence microscopy images. PLoS One. 3, e1997 (2008).

121. S. B. Berman et al., Bcl-xL increases mitochondrial fission, fusion, and biomass in neurons. J. Cell Biol. 184, 707–19 (2009).

122. M. Kislin et al., Reversible disruption of neuronal mitochondria by ischemic and traumatic injury revealed by quantitative two-photon imaging in the neocortex of anesthetized mice. J. Neurosci. 37, 333–348 (2017).

123. M. Arakawa, A scanning electron microscope study of the human glomerulus. Am. J. Pathol. 64, 457–66 (1971).

124. D. Owald et al., A Syd-1 homologue regulates pre- and postsynaptic maturation in Drosophila. J. Cell Biol. 188, 565–579 (2010).

125. Q. Liu, S. Liu, L. Kodama, M. R. Driscoll, M. N. Wu, Two dopaminergic neurons signal to the dorsal fan-shaped body to promote wakefulness in Drosophila. Curr. Biol. 22, 2114–2123 (2012).

126. K. Cichewicz et al., A new brain dopamine-deficient Drosophila and its pharmacological and genetic rescue. Genes, Brain Behav. 16, 394–403 (2017).

127. K. Xu, G. Zhong, X. Zhuang, Actin, spectrin, and associated proteins form a periodic cytoskeletal structure in axons. Science. 339, 452–456 (2013).

